# SuBMIT: A Software Toolkit for Facilitating Simulations of Coarse-Grained Structure-Based Models of Biomolecules

**DOI:** 10.64898/2026.05.18.725912

**Authors:** Digvijay Lalwani Prakash, Arkadeep Banerjee, Shachi Gosavi

**Author notes:** Correspondence to: Shachi Gosavi, Digvijay Lalwani Prakash.

## Abstract

Coarse-grained structure-based models (CG-SBMs; or Gō models) are simplified potential energy functions of biomolecules or biomolecular complexes that encode their structure. Molecular dynamics simulations of such SBMs have been successfully used to study long time-scale dynamics such as protein and RNA folding, and large conformational transitions of biomolecular complexes. SBMs have several advantages: (1) Their MD simulations are computationally inexpensive, making extensive sampling easily accessible to many researchers. (2) They are easy to modify and can be adapted for the specific biomolecular problem that needs to be investigated. However, the force-fields of SBMs are not usually included in commonly used biomolecular simulation packages resulting in a barrier to their use. Here, we present SuBMIT (**S**tr**u**cture **B**ased **M**odels **I**nput **T**oolkit; https://github.com/sglabncbs/submit), a toolkit for generating coarse-grained SBM input files for performing MD simulations with GROMACS and OpenMM/OpenSMOG. Simulations whose input files can be generated using the different flavors of CG-SBMs present in SuBMIT include the folding and conformational ensembles of proteins with intrinsically disordered regions, 3D-domain-swapping in proteins and the dynamics of RNA-protein assemblies (e.g., simple RNA viruses).

## INTRODUCTION

Biopolymers such as proteins and RNA have multi-dimensional folding energy landscapes, where the number of dimensions (degrees of freedom) increases with the length of the polymer. This makes it difficult to extensively sample the landscape to study processes occurring over long length and time scales, such as the folding, binding and assembly of large biomolecular systems. To tackle this, computational models can (a) reduce the degrees of freedom by using a coarse-grained representation of the molecule(s), and/or (b) simplify the interactions that contribute to the potential energy of the model^1–4^. Coarse-grained structure-based models (CG-SBMs), originally invented^5–7^ and successfully used to understand protein folding^8–12^, do both. CG-SBM potential energy functions represent residues using one to a few coarse-grained beads and only native-like interactions (those present in the folded state of the biomolecule) are present between these coarse-grained residues. Although nucleic acid folding landscapes are more frustrated^13^ (include more stabilizing interactions that are not present in the folded state and which can create trapping during folding), CG-SBMs have been used to study the folding of small RNA/DNA molecules^14–16^. Variants of CG-SBMs have also been used to understand protein conformational transitions as well as to simulate the effects of protein-lipid interactions^17–20^ or post-translational modifications^21^ on the folding and conformational transitions of proteins. Finally, CG-SBMs have been used to study protein-protein^22–27^ and protein-nucleic acid^28–30^ binding. This has enabled the studies of both homomeric (fibrillar aggregation^26,31^, 3D-domain-swapping ^9,32–34^) as well as heteromeric^23,25^ assemblies.

Common biomolecular MD simulation packages^35–37^ such as GROMACS^38–40^ or OpenMM^41,42^ are user friendly, broadly accessible and have integrators which can perform fast MD simulations. However, CG-SBM force-fields are not included in these packages despite their obvious utility for understanding the dynamics of biomolecules. MD simulations of CG-SBMs can still be performed using these “standard” MD packages by constructing input files which use in-built functional forms (already encoded in the simulation packages) to generate CG-SBM potential energies. If the required functional form is not present in the MD package, it can be implemented with user-defined or tabulated potentials. But having to generate these CG-SBM input files creates a barrier to the use of these models. Here, we present SuBMIT, a package for generating input files for different flavors of CG-SBMs to perform MD simulations with GROMACS^40^ and OpenMM/OpenSMOG^43^.

The rest of the sections are organized as follows: In the next section, we describe the overall form of CG-SBMs, for which SuBMIT generates input files. We then describe SuBMIT and the details of the implemented models and how they can be tuned using user input. The following section illustrates the use of SuBMIT, with examples which use different models and input flags and arguments. Next, we use SuBMIT to reproduce results from previous studies^44–46^, which were originally simulated using other methods of input file generation. Finally, we compare SuBMIT with other packages which generate SBM input files for MD simulations.

## OVERVIEW OF CG-SBM

Most biomolecular MD packages have in-built atomistic potential energy functions such as CHARMM^47–49^ or AMBER^50–52^ which can then be chosen to perform simulations. These potential energy functions contain different functional forms such as harmonic functions (for bonded terms), cosine functions (to encode periodic angle potentials), Lennard-Jones (LJ)-like functions (to encode breakable attractive interactions and hydrogen bonds), Coulomb interactions (to encode electrostatics), etc. The MD programs are optimized to be computationally efficient at calculating such potential energy terms and their derivatives. CG-SBMs often use the same functional forms as these standard atomistic potential energy functions and can thus be efficiently simulated using the MD simulation packages. In this section, we describe the basic forms of the CG-SBMs. Details pertinent to SuBMIT are provided in the next section. A general form of the CG-SBM potential energy is shown in Equation 1.

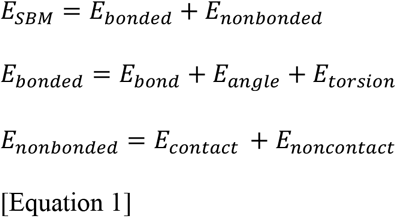

A CG-SBM with the protein coarse-grained to 2 beads per amino acid (a backbone bead: bb or Cα and a side chain bead, sc or Cβ) is implemented as the default model in SuBMIT and is shown below (Equation 1a).

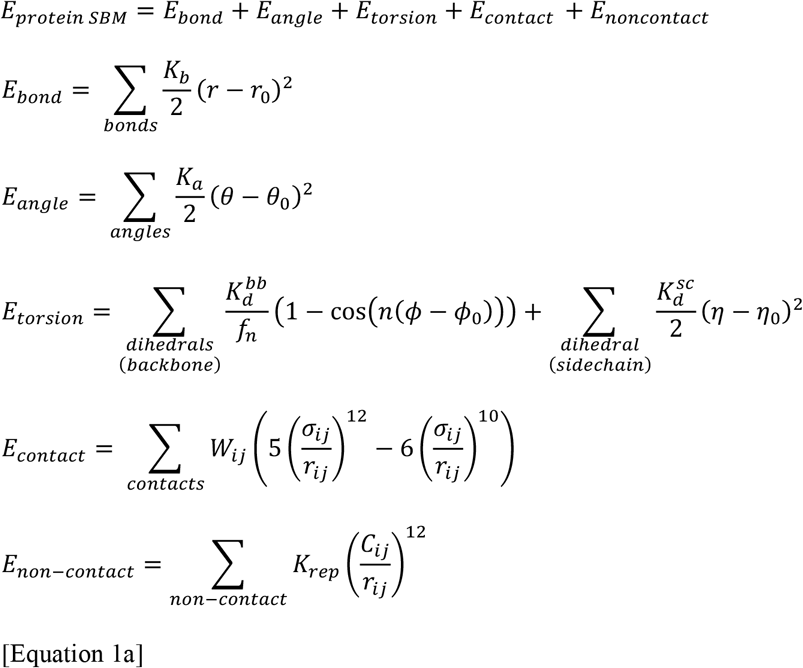

A harmonic function is used to define the potential energy of bond lengths (*E*_*bond*_), angles (*E*_*angle*_) and side-chain dihedrals (the second term of *E*_*torsion*_), while a cosine function (with n=1 and 3 multiplicity; the first term of *E*_*torsion*_) is used for backbone dihedrals. In Equation 1a, *r*_0_, *θ*_0_, *ϕ*_0_ and *η*_0_ are the bond lengths, angles, backbone dihedrals and side-chain dihedrals derived from the native structure while *r, θ, ϕ* and *η* are the bond lengths, angles, backbone dihedrals and side-chain dihedrals calculated from the MD simulation snapshots. Two beads (*i, j*), that do not interact through a bond, angle or torsion potentials, are said to be in contact if they are proximal in the native structure based on some chosen criteria, such as cutoff distance. Since most dynamics occur because of contact formation and breaking, contact calculations are important and detailed in the next section. A LJ-like function is used to define an attractive potential between two beads (*i, j*) in contact, with the native distance between them (*σ*_*ij*_) giving the minimum of the function. For all pairs of beads not forming a contact in the native structure, a repulsive function is defined, where *C*_*ij*_ is the mean of the exclusion volume radii of beads *i* and *j*.

The global minima for all the interactions (except non-contacts) correspond to the parameter values derived from the native structure. Constants 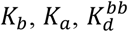 and 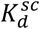 are the force constants for the bond length, bond angle, backbone dihedrals and side chain dihedrals, with *f*_*n*_ being the multiplicity dependent force constant factor. The values of all these force constants can be chosen, but to preserve the chain connectivity and local geometry of the protein, the bond and angle force constants are much larger than the other force constants, whose value is close to that of the basic energy scale of the model, termed ε (in reduced energy units; for both GROMACS and OpenSMOG input file construction, 1ε = 1 kJ mol^-1^).

The nucleic-acids (RNA/DNA) can be coarse-grained to either 1 bead or 3 beads per nucleotide (a backbone phosphate bead: P, a backbone sugar bead: S and a side-chain N-base bead: B). The default CG-SBM implemented in SuBMIT is given below (Equation 1b).

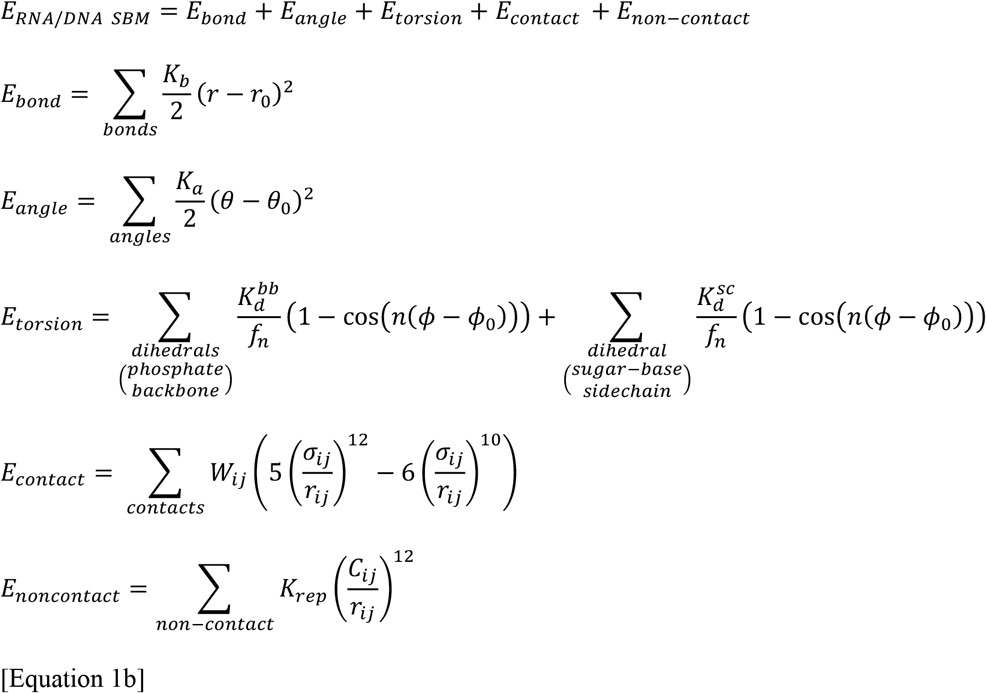

Equation 1a and Equation 1b are similar except for two key differences: (1) Unlike in the protein model (Equation 1a), both the backbone and the sidechain torsions are defined using a periodic cosine function in Equation 1b. The backbone dihedral term is defined between four consecutive phosphate beads (P_i_, P_i+1_, P_i+2_, P_i+3_), while the sidechain dihedral term is defined between the adjacent sugar and N-base beads (B_i_, S_i_, S_i+1_, B_i+1_). (2) Unlike in the protein model, the native contact list in the RNA/DNA model includes beads which interact through a dihedral potential.

However, as in the protein model, RNA/DNA beads that interact through bond or angle potentials are excluded from the contact list. In the default protein and nucleic-acid models, the interaction strengths (or force-constants) have comparable values, allowing the two models to be used together for a protein-RNA/DNA simulation system.

Equations 1a and 1b describe a Cα-Cβ protein and a P-S-B nucleic acid CG-SBM. However, SBMs can differ in the level of coarse-graining and the description of bonded and non-bonded interactions. In addition to the default potentials described above, SuBMIT supports implementation of several SBM variants either by using preset model input flags (Table 1) or through user defined input flags (Table 2; these allow easy modification of default potentials with additional terms). Both these types of flags are discussed in the next section.

**Table 1.**
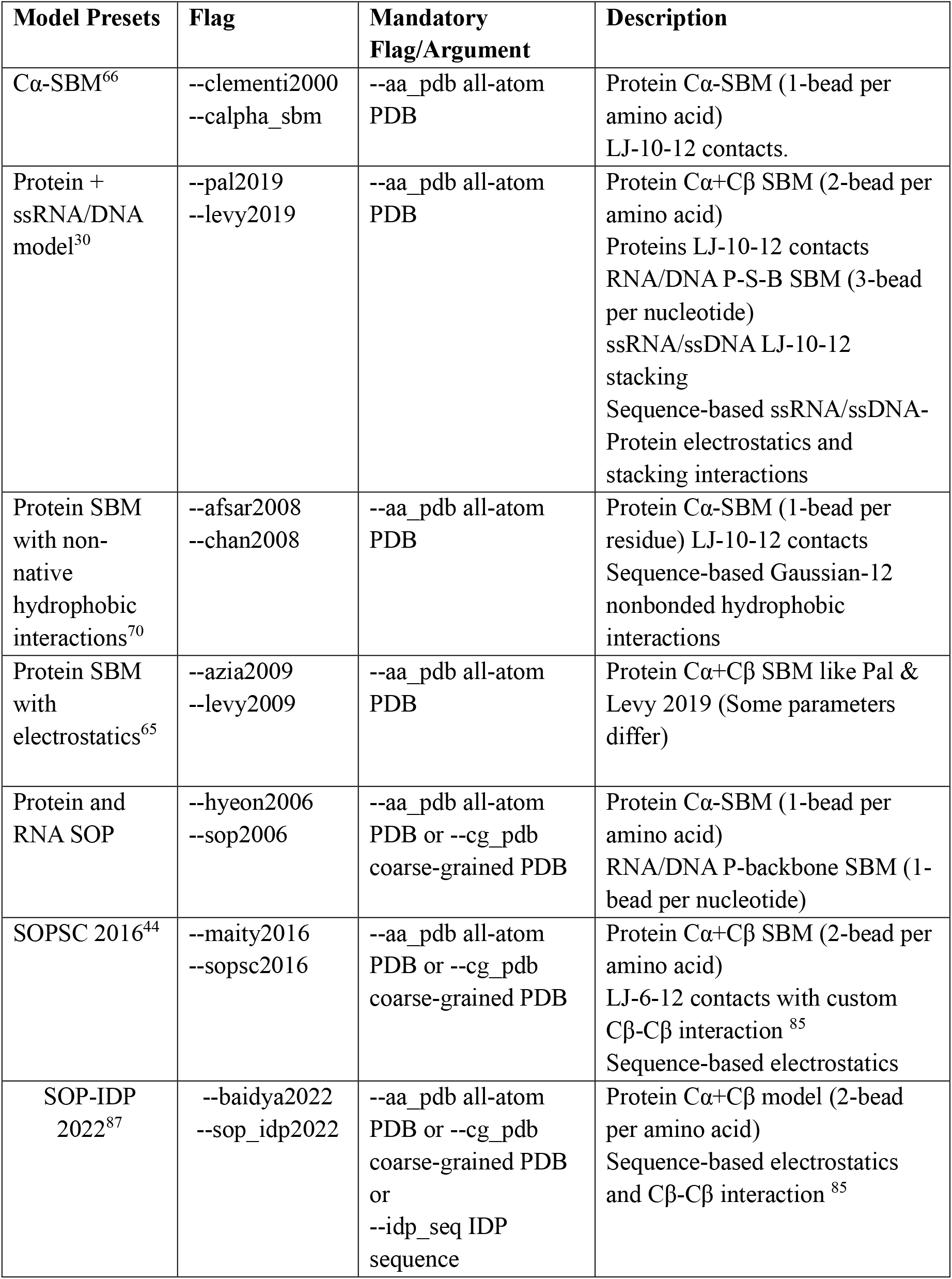
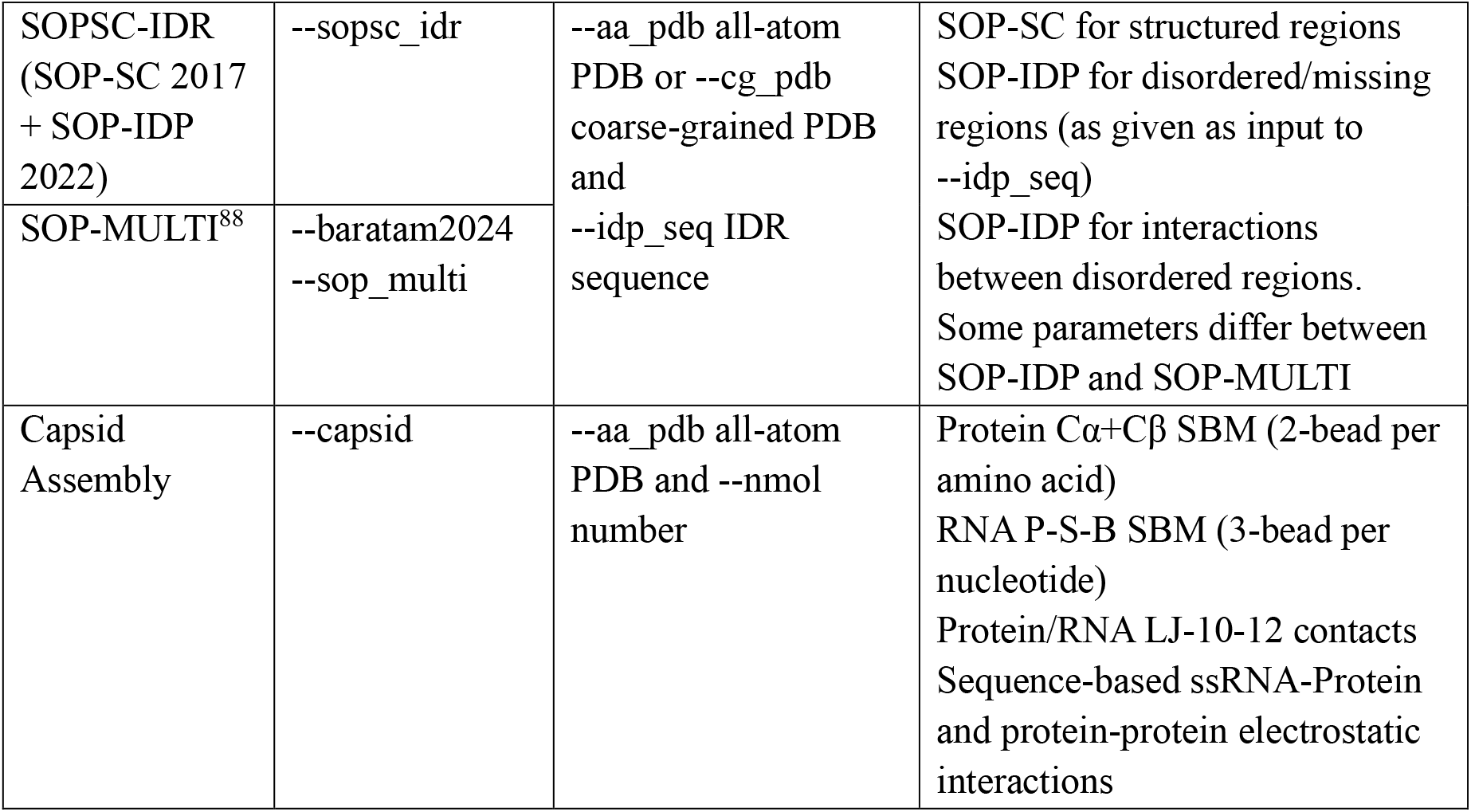
List of preset models included with SuBMIT.

**Table 2.**
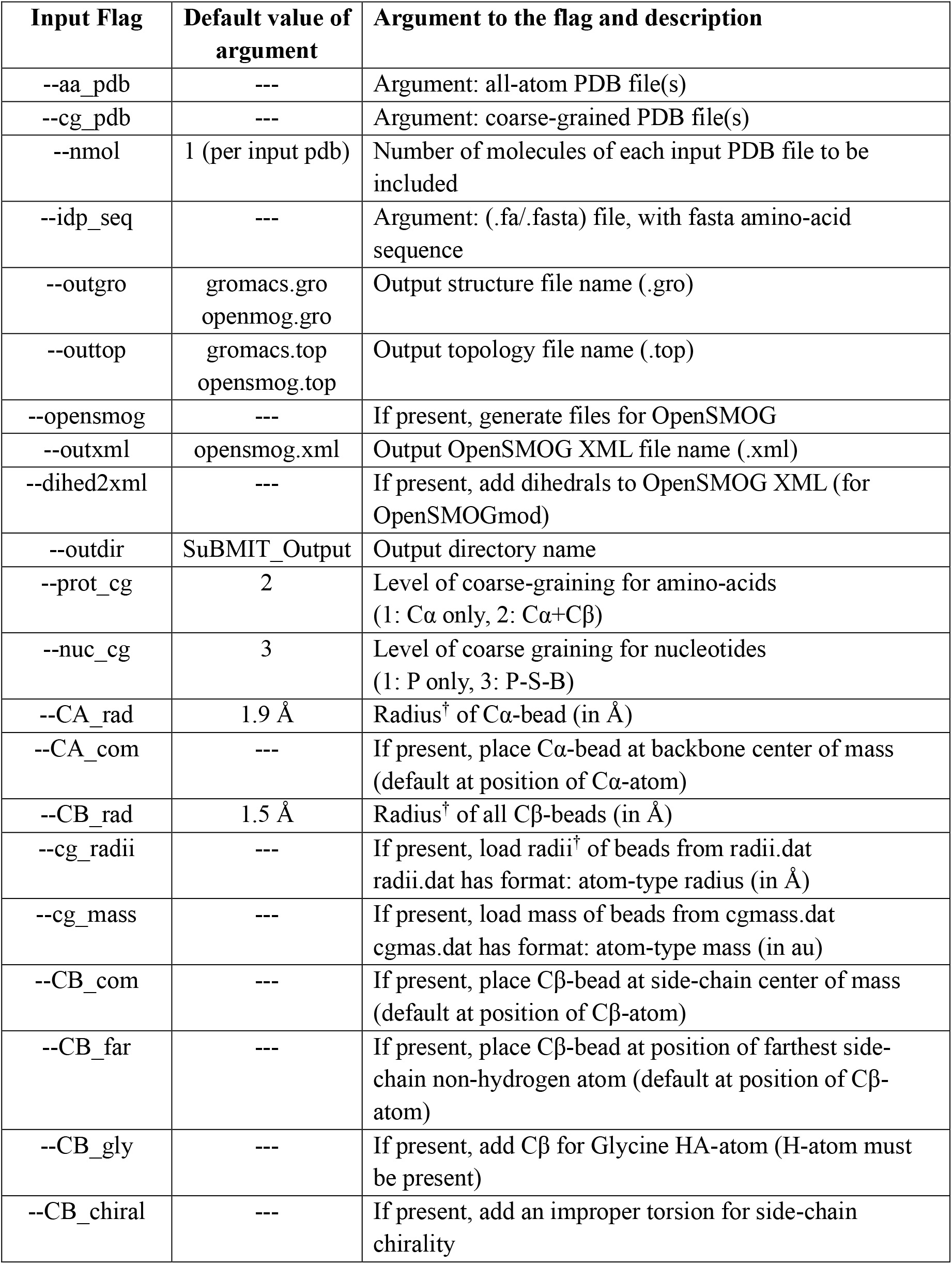
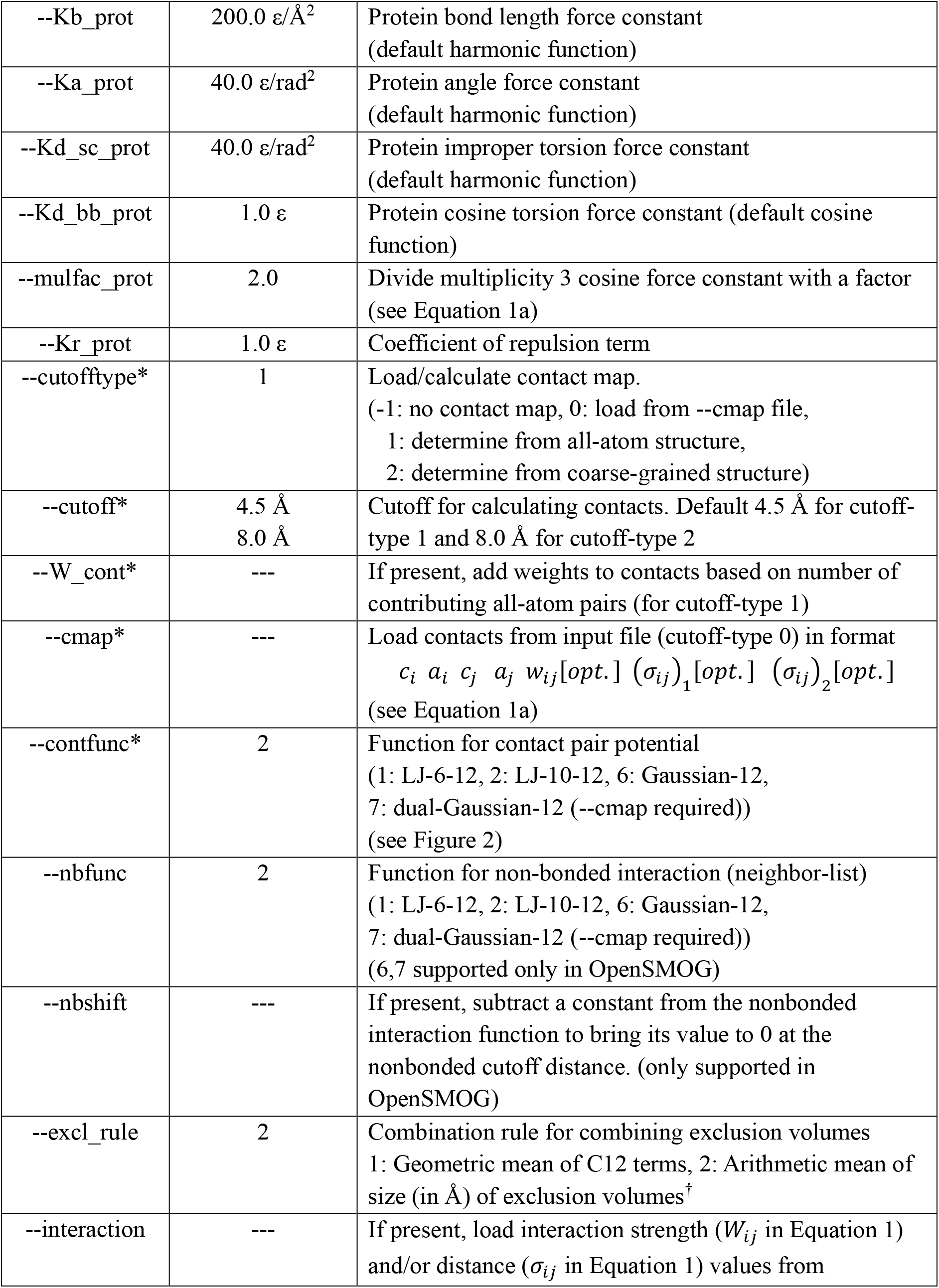
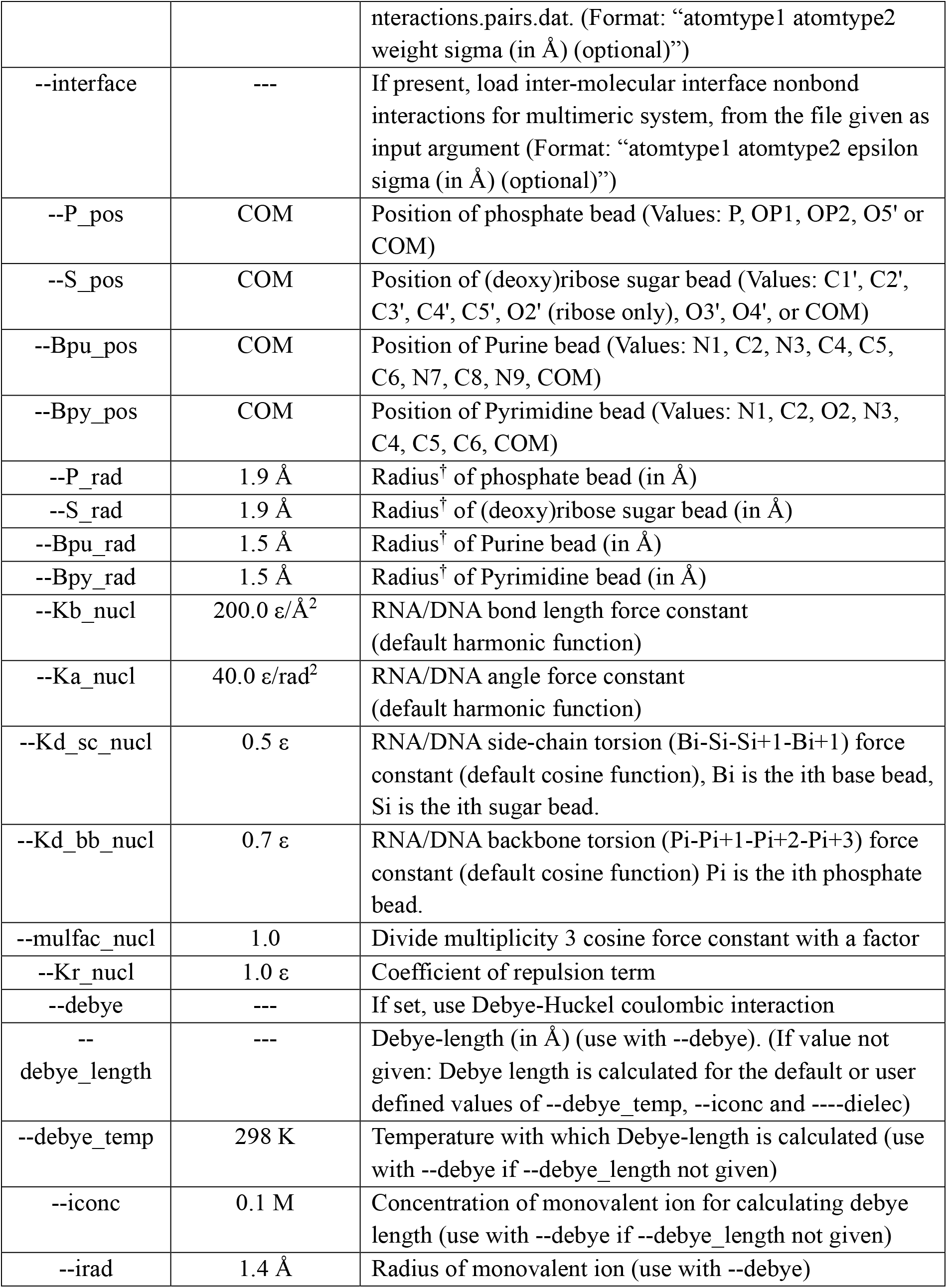
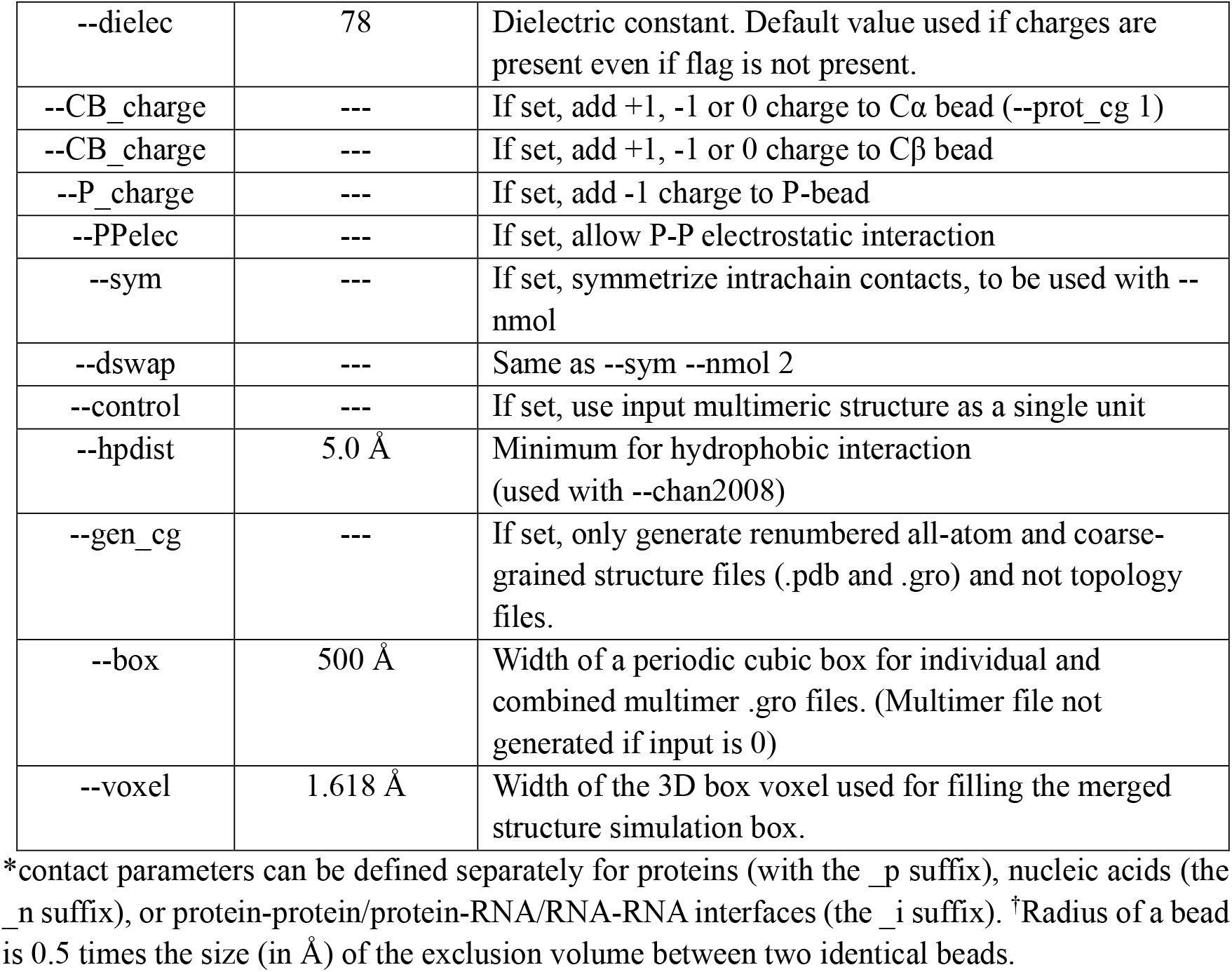
SuBMIT input flags.

## SOFTWARE OVERVIEW

SuBMIT is an open-source Python program, licensed under the GNU Public License version 3. The program can be deployed on systems running Linux, Windows and MacOS with Python 3.10 or above installed. The program takes PDB (.pdb), GRO (.gro) or PDBx/mmCIF (.cif) file(s) with protein and/or nucleic acid structure(s) as input and generates coarse-grained structure (.pdb and .gro) and topology (.top and .xml) files, which encode the structure-based potential. These files can then be used to perform molecular dynamics simulations on either GROMACS (version 4.5; .gro, .top)^40^ or OpenSMOG (version 1.x; .gro, .top, .xml)^43^ which is based on the OpenMM Python API^41,42^.

To define the structure-based potential, SuBMIT accepts multiple input flags in addition to the structure file. Examples of few such parameters include level of coarse-graining, values of force constants, type of non-bonded interactions, number of molecules, additional sequence-based interactions, etc. Moreover, the program includes multiple preset models with predefined input parameters that can be directly invoked using a single flag: the name of the model. Compatible modifications to the predefined parameters of any given model can be made by using the optional input flags. The list of currently supported preset models (Table 1) and the list of supported input flags with their default values (Table 2) can be viewed by using the --help flag.

### SuBMIT architecture and workflow

Figure 1 shows SuBMIT’s workflow. The package consists of three components: submit.py, PDB_IO.py and topology.py. The submit.py handles user-input and default flags for different models and is the main script run by the user (e.g., python ${SuBMITDIR}/submit.py, where ${SuBMITDIR} is the path of the repository directory). The PDB_IO.py includes classes to handle reading, loading, cleaning, renumbering and writing all-atom and coarse-grained PDB (.pdb) and GRO (.gro) files. To support mmCIF input files (usually used for larger structures) PDB_IO converts mmCIF to PDB files, with systems with >=10^6^ atoms being converted to PDB with hybrid-36 based numbering^53^. Originally developed and included with the later versions of CCBTX-package (computational crystallography toolbox)^54^, the hybrid-36 format is a common numbering method also used by tools such as MODELLER^55^ and UCSF-Chimera^56^ for writing large PDB files. For each input file, PDB_IO creates a protein and/or nucleic acid structure-data object(s), which holds information from the input files, and this is passed onto topology.py. The topology.py includes a set of classes for determining structure-based parameters from the input structure-data objects and these parameters are written to the topology file (.top for GROMACS; .top and .xml for OpenSMOG).

**Figure 1.**
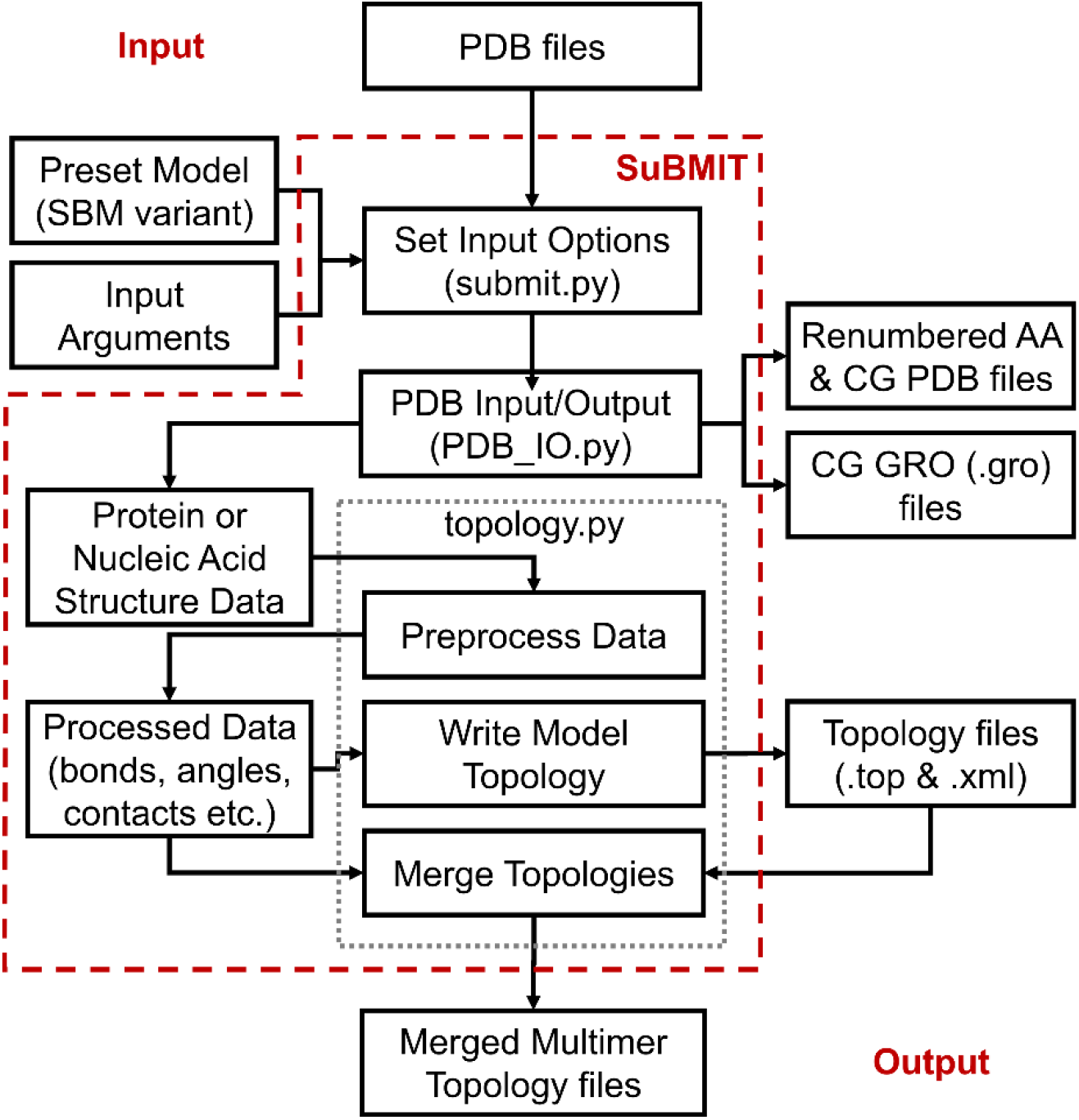
SuBMIT design and workflow showing components, inputs and outputs.

### Setting the level of coarse-graining

SuBMIT supports two levels of coarse graining each for proteins (1 or 2 beads per residue; set using the flag --prot_cg) and nucleic acids (1 or 3 beads per nucleotide; set using the flag --nucl_cg) (Table 2). For proteins, in the 1-bead representation, the whole residue is represented by a single bead, generally placed at the position of the Cα atom (Cα-bead). In the 2-bead representation, the whole residue is represented by two beads: a backbone bead (Cα) and a second sidechain bead (Cβ), placed at the position of the Cβ atom, or at the center of mass of the sidechain, or at the position of the farthest heavy (non-hydrogen) atom of the sidechain. For RNA/DNA structures, in the 1-bead representation, a single bead (P-bead) is placed at the position of the backbone phosphate (P) atom or at the center of mass of the phosphate group. In the 3-bead representation, along with the backbone phosphate P-bead, additional beads representing the (deoxy-)ribose sugar (S-bead) and the N-base (B-bead) are used. These beads can either be placed at the center of mass or at the position of a specific atom in the given group.

### Contact-maps

In most common variants of SBMs, a non-bonded attractive potential is only defined between two beads which are not part of a bonded interaction (bond, angle or dihedral) and which are proximal (or in contact) in the native structure. For a structure with N coarse-grained beads, the contact map is an N-by-N binary matrix whose (*i, j*) and (*j, i*) terms are 1 if beads *i* and *j* are in contact in the native structure and 0 otherwise. Contact maps (or proximity between beads) have been calculated in multiple ways such as by using the Shadow Contact Map (SCM) algorithm^57^, the Contacts of Structural Units (CSU) algorithm^58^ or by using a simple cutoff algorithm^59^. SuBMIT takes a --cutofftype flag to determine which contact map to calculate and/or use. When --cutofftype 1, the contact map is calculated using the all-heavy atom structure using a predetermined cutoff distance (default 4.5 Å) and then mapped back onto the coarse-grained representation. So, two beads are said to be in contact if any of their corresponding heavy (non-hydrogen) atoms are within a cutoff distance of each other. Thus, multiple all-atom contact pairs can contribute to a given contact between the corresponding CG-beads. This “weight” can either be ignored (set to 1 for all contacts resulting in a binary contact map; this is the default) or used to create a weighted contact map, each element of which is either 0 (no contact) or a weight, *W*_*ij*_ (Equation 2). To maintain the relative contribution of the nonbonded and bonded interactions in the potential energy function, the total potential energy of the weighted contacts is scaled such that it is comparable to the potential energy contribution of the unweighted contacts^60–62^ as follows,

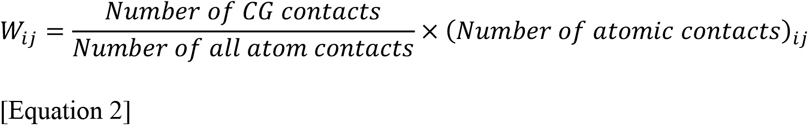

Here, (*Number of atomic contacts*)_*ij*_ is the number of atomic contacts between beads (*i, j*). A weighted contact map can be implemented using the --W_cont flag.

A contact map can also be calculated directly from the coarse-grained representation using -- cutofftype 2. With this argument, a contact is present if two coarse-grained beads are within a cutoff distance (default 8.0 Å). Weights are not used with this argument. The --cutofftype flag can also take an input of -1, when no contact map calculation is to be performed. Finally, --cutofftype 0 can be used to upload a user-defined contact map file. This allows the use of custom weights or uploading screened contact maps such as SCM or CSU maps. The user-defined contact map file can be given using the --cmap [filename] flag, in the format:

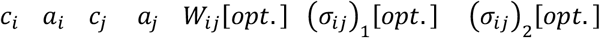

Here, each line specifies a contact between coarse-grained beads *i* and *j* with (*c*_*x*_, *a*_*x*_) being the chain identifier and the atom number for bead *x*. The *W*_*ij*_ (user-defined weight) and *σ*_*ij*_ (native distance of contact in Å) are optional parameters. The contact-map file can include up to two distance values, where the second value is only used for the two minimum Gaussian potential (see next subsection).

The contact map parameters (e.g., cutoff) can be defined separately for proteins, nucleic acids and biomolecular (protein/RNA/DNA) interfaces using the “_p”, “_n” or “_i” suffixes with the corresponding input flag (e.g., --cutofftype_n for RNA/DNA; see Table 2). By default, the interface contact potential between molecules input through separate PDB files is not calculated (--cutofftype_i -1). A user-defined contact map file with weights and distances has to be provided for such systems.

### Non-bonded contact potential

SuBMIT supports different forms of bonded and non-bonded potential functions. Equation 1a shows a representative potential energy function of a two bead protein CG-SBM, which is the default implementation in SuBMIT. In SBMs, a contact potential is the function used to encode an attractive potential between the beads in contact. Since Lennard-Jones (LJ)-like nonbonded potentials are implemented (and their computation is optimized) in most biomolecular MD simulation packages, the same functional form is often also used to encode contact potentials in SBMs^60,63–69^ . An example of a LJ-like function is present in Equation 1a. SuBMIT supports encoding two different LJ-like functions which can be selected using the contact function (--contfunc) flag. These are

i. LJ-6-12 (--contfunc 1):

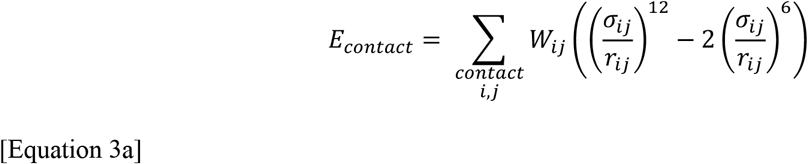
ii. LJ-10-12 (--contfunc 2):

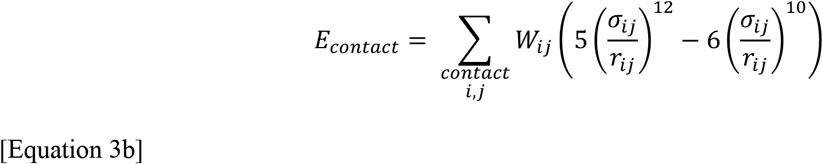

*σ*_*ij*_ and *r*_*ij*_ are the distances between the beads *i* and *j* in the native structure and the MD snapshot, respectively. *W*_*ij*_ is the strength of the contact potential. In both the expressions, the 12-power term is the repulsive component of the function and dominates when *r*_*ij*_ < *σ*_*ij*_. The 6-power and the 10-power terms are the attractive components of the functions and dominate when *r*_*ij*_ > *σ*_*ij*_. The minimum of the LJ-function is at *r*_*ij*_ = *σ*_*ij*_. Since the 10-power attractive term decays faster to zero, using it makes for a narrower attractive minimum. This is preferred over the 6-power term for CG-models^64–66,68^, where the *σ*_*ij*_ are larger than in atomistic SBMs.

LJ-like functions are limited because the “contact” distance also determines the size of the beads (the excluded volume or the repulsive interaction). Contact distances are large in coarse-grained interactions, and this can lead to large energies when all attractive interactions are not consistent with each other. This can occur when encoding non-native attractive interactions (such as a general hydrophobic attraction^70^) or the multiple interconverting structures of the same sequence^71^. Additionally, it is difficult to analytically implement two minima (say for a conformational transition^20^) in LJ-like potentials. The Gaussian contact potential^72^ overcomes these limitations^70,72–75^ by having one distance to define the repulsive term and a different distance to define the attractive term. Additionally, multiple inverted Gaussian functions can be used to define multiple attractive minima. SuBMIT can be used to encode both a single minimum (G-12) and a dual-minima (G1-G2-12) Gaussian contact potential:

i. G-12 (--contfunc 6):

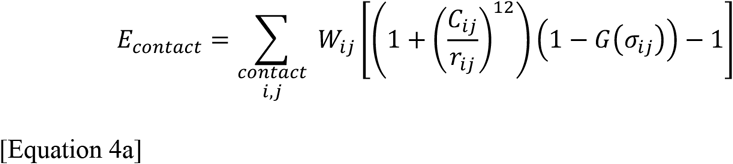

where the *C*_*ij*_ in the repulsive 12-power term sets the radius of the excluded volume between beads *i* and *j*. The Gaussian term *G* is given by 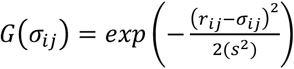, with *σ*_*ij*_ and *r*_*ij*_ being the distances between the beads *i* and *j* in the native structure and the MD snapshot, respectively. *σ*_*ij*_ sets the maximum of the Gaussian term and therefore, the minimum of the potential. The width of the Gaussian function is set by defining the standard deviation, *s* (= 0.5 Å).
ii. G1-G2-12 (--contfunc 7):

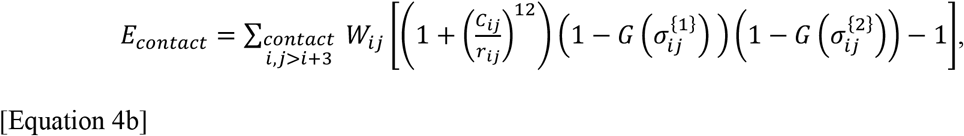

Equation 4b shows the expression for a potential which encodes two Gaussian minima. In this dual-Gaussian, 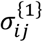 and 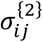 are the two minima of the contact potential. Figure 2 shows a comparison of the above 4 contact potentials (Equations 3a, 3b, 4a and 4b).

**Figure 2.**
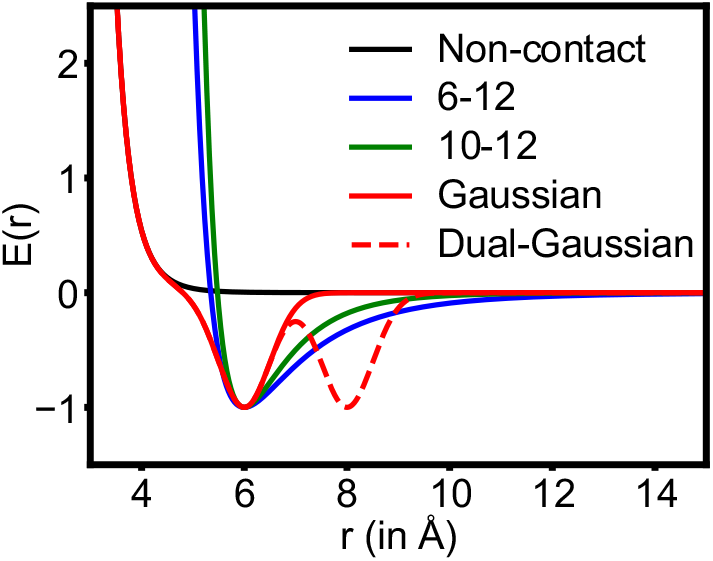
Potential energy E(r) versus distance (r) plots for different nonbonded interaction functions supported by SuBMIT. Non-contact beads interact through a purely repulsive interaction term (*E*(*r*) = *E*_*non*−*contact*_ in Equation 1a & 1b; black). For beads with attractive interactions (*E*(*r*) = *E*_*contact*_, in Equation 1a and 1b) SuBMIT supports Lennard-Jones (LJ) like functions, LJ-6-12 (Equation 3a; blue), LJ-10-12 (Equation 3b; green). The program also supports a combination of a repulsion term with a single negative Gaussian function for a single minimum (Equation 4a; red solid line) or a combination with two negative Gaussians for a two minima function (Equation 4b; red dashed line). The native distance (*σ*_*ij*_ between beads *i* and *j*) for the single minimum attractive potential (Equations 3a, 3b and 4a) is *σ*_*ij*_=6.0 Å. There are two native distances (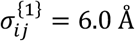 and 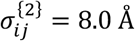) for the two minimum attractive potential (Equation 4b). The width of the Gaussian functions (Equation 4c) is *s*=0.5 Å. The excluded volume term between beads *i* and *j, C*_*ij*_ is set to 3.8 Å (in Equation 1, 4a and 4b). The weights (prefactors) of all functions are set to 1.

It is important to note that by default, GROMACS only supports the LJ-6-12 function. For LJ-10-12, SuBMIT automatically generates GROMACS supported tabulated pair potentials in “table files” which can be used with the GROMACS mdrun command. The Gaussian potential can only be implemented on a modified version of GROMACS 4.5.4 which is available at the SMOG-server (https://smog-server.org/extension/#gauss). In contrast, all the above-mentioned functions can be directly implemented on OpenSMOG^43^.

In addition to LJ and Gaussian contact potentials, the “desolvation-barrier” (DSB) contact potential^76–78^ is also used in some SBM variants. The DSB contact potential is similar to the LJ potential but adds a shoulder/barrier at a distance (equal to the radius of the water molecule) from the native contact distance (*σ*_*ij*_). The barrier accounts for the cost associated with the expulsion of a water molecule, when the native contact is formed. While not included in the current version, support for the DSB contact potential will be added in future versions of SuBMIT.

### Symmetrized and equivalent contacts

SuBMIT can handle biomolecular complexes which have multiple copies of a given biomolecule. This is done by creating the required number of copies of the molecule and duplicating the contacts. Equivalent contacts are usually interface contacts between two molecule types and are defined for all possible combinations of the molecule pair. For instance, if a system has two copies of molecule A (A1, A2) and three copies of molecule B (B1, B2, B3) and if A interacts with B, then for each contact pair A_*i*_–B_*j*_, the program adds 6 entries corresponding to A1_*i*_–B1_*j*_, A1_*i*_–B2_*j*_, A1_*i*_–B3_*j*_, A2_*i*_–B1_*j*_, A2_*i*_–B2_*j*_ and A2_*i*_–B3_*j*_ to the contact list. So, for a system with N_A_ molecules of A and N_B_ molecules of B, N_A_ x N_B_ equivalent copies of each A-B contact get added to the contact list. Equivalent contacts were recently used in the GoCa-model^25^ to study the assembly of homo-multimeric systems.

In contrast to equivalent contacts, which are copies of interface contacts, symmetrized contacts are used when contacts can form in either an intrachain or an interchain fashion between a pair of identical molecules^9,32,34^. For instance, if a system has 2 copies (A1 and A2) of a given molecule A and a contact (*i, j*) exists in the native structure of A, then the program will add 4 entries for the contact, corresponding to A1_*i*_–A1_*j*_, A1_*i*_–A2_*j*_, A2_*i*_–A1_*j*_, A2_*i*_–A2_*j*_, to the contact list. So, for a system with N-copies of a molecule, the program will add N^2^ symmetrized copies of each intra-molecular contact to the contact list. Example systems where symmetrized contacts are used in simulations are the folding and assembly of domain swapped proteins^32–34,45,79,80^ and the folding of proteins in the presence of their own fragments^46,81^.

For both equivalent and symmetrized contacts, the number of contacts to be defined in the MD topology file increases by a power of 2 (N^2^), when N is the number of molecules to be simulated. For larger biomolecular assemblies, this leads to a large topology file. Additionally, the atom number dependent pair potential entries in both GROMACS and OpenSMOG are handled with a static pair list, similar to the bond-length potential. So, the potential is calculated for all the included atom pairs, irrespective of the distance between the beads in the simulation box. This reduces computational efficiency in large simulations. To handle this, for systems with more than five copies of a molecule, SuBMIT creates a separate unique atom type for each bead involved in a symmetrized or an equivalent contact. The corresponding atom-type pair is then added to the nonbonded parameter section of the topology file. Unlike directly defined atom number pairs (in the GROMACS pairs section or OpenSMOG contacts section), nonbonded parameters in both GROMACS and OpenSMOG are handled using the neighbor list algorithms^82^, making computation faster.

### Preset Models

There are several predefined models present in SuBMIT (Table 1). These have specific settings for level of coarse-graining, value of force constants, strength of non-bonded interactions, choice of sequence-based interactions, etc. Some preset models require non-standard types of bonded or non-bonded functional forms that are not part of the default implementation. These are hard-coded as part of the model and are not directly available for use with new models (see next subsection). The included preset models are of the following types: (1) <u>Purely structure-based models:</u> only information derived from the input structure is used to encode the potential energy function, with force constants and non-bonded interaction strengths predefined as part of the model (for instance, a commonly used Cα-SBM^66^). It should be noted that some models have contacts derived only from the structure but the strength (or the weight) of the interaction is determined by the identity of the residues which are in contact. Such weights can be statistically derived from known protein structures or from experimental studies (knowledge-based^1,2^). Example of such weights include the Miyazawa−Jernighan^83,84^ and the Betancourt-Thirumalai^85^ pair potential for proteins. The latter is used by the self-organized polymer (SOP) class of models^44,67,69,86–88^. (2) <u>CG-SBMs that have some additional structure-independent (potentially sequence based) interactions:</u> Examples include a Cα-SBM with hydrophobic interactions^70^, SBMs with sequence-dependent electrostatic interactions^30,44,65,67,87^ and SBMs with purely sequence-based interactions for intrinsically disordered proteins^87^ or regions^88^. For nuclei acids, SuBMIT supports a model with a custom base-stacking potential^29,30^, which is defined between two nucleic acid N-bases or between a nucleic acid N-base and protein aromatic sidechains. It also supports an SBM with electrostatics for both proteins and single stranded nucleic acids to study protein-ssDNA/ssRNA binding^29,30^. Finally, SuBMIT also has presets which help use different variants of SBMs for studying multimeric folding (such as domain swapping^32–34,45,79,80^, using the flag: -- dswap) or assembly (such as virus-assembly, using the flag: --capsid).

### Implementing new models with SuBMIT

We expect that input files for most new models can be created by choosing appropriate input flags and arguments (Table 2). However, some models might require a significant modification of one or more functions. In such cases, a new child class can be created in topology.py (or a different python file) which inherits the Topology class. The Topology class (present in topology.py) includes a set of functions, each aimed at writing a particular section in the GROMACS (.top) and OpenSMOG (.top and .xml) topology files. Depending on the features required in the new model, specific methods can be re-written in the child class without having to modify the Topology class. While running the program with the new model input flags, the new methods will override the ones in the Topology class, while all other methods will work as they are. This modularity allows for the easy addition and testing of new models, which can later be integrated into SuBMIT if required. Implementation of new models is detailed on the wiki section of the SuBMIT repository (https://github.com/sglabncbs/submit/wiki).

## SuBMIT USAGE ILLUSTRATED WITH EXAMPLES

SuBMIT can be used either by providing multiple input flags or by choosing preset models. A list of available preset models (Table 1) and a list of optional input flags (Table 2), can be generated using the help flag (--help):

~~~
                  python submit.py --help
~~~

[Example 1]

We first describe some example use cases of SuBMIT which utilize input flags (Table 2). All examples are included with the program repository.

### Generating files for a single input structure

The PDB file can be given as an argument to the --aa_pdb flag for generating input files for small proteins, like ubiquitin (PDB: 1UBQ), using the default SuBMIT parameters:

~~~
                  python submit.py --aa_pdb 1UBQ.pdb
~~~

[Example 2]

The all-atom input PDB flag (--aa_pdb), is a mandatory flag for using the default parameters. The output files include a “gromacs.gro” and a “gromacs.top” file. The “gromacs.gro” contains the coarse-grained 2-bead structure of the protein. The “gromacs.top” contains sections corresponding to bonded and non-bonded interactions (Equation 1a). Flags --outgro and --outtop can be used to override the default output file names. The default LJ-10-12 pairs function can only be implemented using tabulated pair potentials for which the table file “table_lj1012.xvg” is also generated. Additionally, the program also writes a contact map file “prot.CGcont”. By default, the contact map is calculated using the all-atom representation of the structure with a 4.5 Å cutoff. Hence, an additional “prot.AAcont” is also generated. The OpenSMOG flag (--opensmog) can be used to generate input files for OpenSMOG.

~~~
                  python submit.py --aa_pdb 1UBQ.pdb --opensmog
~~~

[Example 3]

The Example 3 code generates “opensmog.gro”, “opensmog.top” and “opensmog.xml”. The .top file includes the atom type definitions and bonded interactions, and the .xml file includes the nonbonded interactions and charges. No table file is generated when using the OpenSMOG flag.

The “--CB_charge” flag can be used to add charges to positively (+1) or negatively (-1) charged amino-acid sidechains. When charges are present, the generated output files are different in the following ways: (a) Cβ beads have +1, -1 or 0 charge in the atoms section of “gromacs.top” and (b) an additional table file “table_coul_lj1012.xvg” is generated. The “--debye” flag can be used to switch to the Debye-Hückel potential instead of regular Coulomb electrostatics.

~~~
                  python submit.py --aa_pdb 1UBQ.pdb --CB_charge --debye
~~~

[Example 4]

The Debye-Hückel potential can be tuned using additional parameters such as monovalent ion concentration (--iconc), temperature (--debye_temp), etc. The --P_charge flag for nucleotides is similar to the --CB_charge for proteins and can be used to add a -1 charge to the phosphate bead (P).

For systems with both proteins and nucleic acids, along with the default output files, the program generates separate protein and nucleic acid output files with the “prot_” and “nucl_” prefixes, respectively. For example, for CDC13 DNA binding domain in complex with 11-mer ssDNA (PDB: 1S40), the command

~~~
                  python submit.py --aa_pdb 1S40.pdb --CB_charge --P_charge --debye
~~~

[Example 5]

generates “gromacs.gro” and “gromacs.top”. Additionally, the command will also generate “prot_gromacs.gro”, “prot_gromacs.top” for the protein structure and “nucl_gromacs.gro”, “nucl_gromacs.top” for the ssDNA structure.

### Generating files for multimeric systems

SuBMIT can be used to generate topology files for simulating multimeric systems using either symmetrized or equivalent contact interactions. For example, the following command can be used to study domain swapping of ubiquitin (1UBQ) using a Cα (1-bead per amino-acid) SBM.

~~~
                  python submit.py --prot_cg 1 --aa_pdb 1UBQ.pdb --dswap
~~~

[Example 6]

The domain-swapping flag (--dswap) tells the program to have two identical copies of the protein and symmetrize contacts. The behavior of --dswap can be replicated with --sym --nmol 2. The -- sym flag can also be used with more than 2 molecules:

~~~
            python submit.py --prot_cg 1 --aa_pdb 1UBQ.pdb --sym --nmol 7
~~~

[Example 7]

This symmetrizes intra-chain contacts (determined from the monomer structure) across 7 molecules of ubiquitin. Additionally, interface contacts between all copies of the chains can be replicated for a homo-multimeric system by calculating equivalent contacts. Since the interface contacts cannot be calculated from a single chain, these are provided separately using the --cmap_i flag. For example, to generate files for simulating the assembly of coat proteins of the satellite tobacco mosaic virus (STMV), SuBMIT uses a single coat protein (the asymmetric unit, PDB 1A34) as an input along with an interface contact map file.

~~~
       python submit.py --aa_pdb 1A34.pdb --nmol 60 --cmap_i inter.contacts
~~~

[Example 8]

The file “inter.contacts” contains all the interface contacts of one coat-protein monomer (PDB 1A34 asymmetric unit) with all its neighboring coat-proteins in the virus structure. If the --cmap_i flag is used, the interface contact cutoff type (--cutofftype_i) is automatically set to 0. Additionally, with --cmap_i given, if the number of molecules (--nmol) is more than 1, then the set of interface contacts are automatically duplicated for all equivalent molecule pairs.

For generating files for heteromeric systems, multiple protein and nucleic acid structures can be given as multiple inputs to the --aa_pdb flag. For each input structure (considered as a single molecular unit), the program generates individual output structure and topology files with either “protX_” or “nuclX_” prefix, where X={0,1,2,..} is the index based on the order of inputs to the --aa_pdb flag. An example of a hetero-multimeric assembly system, is the phage Qβ with coat protein dimers (referred to as AB and CC dimers) assembled around its RNA genome. For structure-based RNA-protein and protein-protein interface contacts, and sequence-based RNA-protein and protein-protein electrostatic interactions, the following command can be used:

~~~
     python submit.py --aa_pdb AB.pdb CC.pdb gRNA.pdb --nmol 60 30 1
     --CB_charge --P_charge --debye --cmap_i inter.contacts --opensmog
~~~

[Example 9a]

The --nmol flag is used to instruct the program to create a topology file consisting of 60 copies of input 1 (AB.pdb, AB dimer), 30 copies of input 2 (CC.pdb, CC dimer) and a single copy for input 3 (gRNA.pdb, genomic RNA). If --nmol is not given, a single copy of each input is used. Similar to the homo-multimeric system, interface contacts are not determined for a hetero-multimeric system by default and therefore, are not added to the topology file. In Example 9a, interface contacts are provided separately using the --cmap_i flag. As stated previously, when multiple copies of a given molecule are present, the program automatically generates equivalent interface contacts for all the copies. The interface contact map file (“inter.contacts” in the above example command) contains all protein-protein (AB-AB, AB-CC, CC-AB) and protein-RNA contacts (AB-RNA, CC-RNA) obtained from all the unique interfaces present in the whole capsid structure (PDB: 7LHD). Unlike for the intrachain contact map files, the weight and the distance columns are mandatory in the interface contact map file. Additionally, the chain identifiers must also include prefix “protX_” or “nuclX_” for identifying the input PDB to which the chain belongs. In the above example, prefixes “prot0_”, “prot1_” and “nucl2_” were used for chains corresponding to input 1 (AB dimer), input 2 (CC dimer) and input 3 (genomic RNA), respectively. The prefixes are the same as the ones assigned to the coarse-grained structure and topology files generated by the program for the individual input PDB files. Below are some example lines from the “inter.contacts” file (in format *c*_*i*_ *a*_*i*_ *c*_*j*_ *a*_*j*_ *W*_*ij*_ *σ*_*ij*_).

~~~
             prot0_1 1    prot0_2   301   0.6     8.596
             prot0_1 4    prot1_1     4   2.4     5.455
             nucl2_1 8    prot0_2   366   1.2     7.568
             nucl2_1 224  prot1_2   112   1.8     10.468
~~~

[Example 9b]

In addition to the inter-molecular contact file whose use is shown in Example 9, user-defined intra-molecular contact map files can also be provided as input, instead of being calculated by the program as shown in previous examples. However, since the program reassigns atom numbers for every input structure, it is recommended that it be first run with the “--gen_cg” flag to generate renumbered all-atom and coarse-grained structure files, without generating the topology files as follows:

~~~
      python submit.py --aa_pdb AB.pdb CC.pdb gRNA.pdb --gen_cg
~~~

[Example 10a]

This command generates all-atom PDB files “AB.renum.pdb”, “CC.renum.pdb” and “gRNA.renum.pdb” with reassigned atom numbers. It also generates coarse-grained structure files “prot0_gromacs.gro”, “prot1_gromasc.gro” and “nucl2_gromacs.gro”. These files can be used to generate custom contact maps, which can then be given as input to SuBMIT with the order of the input contact map files same as the order of their corresponding input all-atom PDB files.

~~~
      python submit.py --aa_pdb AB.renum.pdb CC.renum.pdb
      gRNA.renum.pdb --debye --CB_charge --P_charge --nmol 60 30 1
      --cmap_i inter.contacts --cmap AB.txt CC.txt gRNA.txt
~~~

[Example 10b]

#### Merged output files

Along with the topology and structure files for the individual inputs, the commands in Examples 8, 9 and 10, also generate merged structure (gromacs.gro/opensmog.gro) and topology files (gromacs.top for GROMACS or opensmog.top and opensmog.xml for OpenSMOG). In the merged structure file, coordinates corresponding to the copies of the first input to the --aa_pdb flag, are added first, followed by entries corresponding to the copies of second input and so on. The simulation box is filled using a grid-based algorithm, where the cubic box (with edge length defined using --box flag, default 500 Å) is divided into multiple cubic cells/voxels (with edge length defined using the --voxel flag, default 1.6 Å). Each input structure copy is allocated a cubic volume consisting of the minimum number of voxels required to completely fit the given structure, irrespective of its orientation. The first copy of the largest input structure is always positioned in the box center, while all the other copies of all the input structures are randomly rotated and translated in the box ensuring that the cubic volume of each structure does not overlap with the volume of other structures. If the program is unable to create a merged structure file for the given box (--box) and voxel (--voxel) edge-lengths, then it produces a shell script that has a series of GROMACS genbox^40^ commands required to generate the merged structure file. Similar to the structure file, for each section (atoms, bonds, angles, pairs, etc.) in the merged topology files (.top and .xml), entries corresponding to the first argument to the --aa_pdb flag are added first, followed by the copies of other inputs in the same order. For less than five total copies of the input structures, the interface contacts are duplicated and added to the contact list as part of the OpenSMOG contact potential (or GROMACS pair potential). For systems with more than five copies of the input structure, instead of duplicating the atoms for all the copies, each interface “contacting” atom is assigned a unique atom-type and the corresponding contact pair is defined as the atom type pair in the nonbonded parameters section. This is more efficiently handled by the neighbor list algorithm^82^.

### Using SuBMIT with preset models

SuBMIT comes conveniently pre-loaded with multiple preset models (Table 1) that can be used without providing individual flags. For example, to simulate the CDC13 DNA binding domain (PDB: 1S40) with a previously developed model^30^ used to simulate single-stranded-nucleic acid-protein binding, the following command can be used.

~~~
             python submit.py --pal2019 --aa_pdb 1S40.pdb
~~~

[Example 11]

This model uses a 2-bead representation for amino-acids with a structure-based potential and a 3-bead representation for nucleotides. The model encodes a non-structure-based potential for single-stranded (ss)RNA/ssDNA, while the protein-nucleic acid interface interactions are defined using sequence-based Coulombic and stacking interactions.

For every preset model, some parameters can be overridden by using flags. For example, the ssDNA angle force constant can be changed to 80 ε/rad^2^ (from the default = 40 ε/rad^2^; as stated earlier, ε is set to 1 kJ mol^-1^) by using the --Ka_nucl flag as follows:

~~~
       python submit.py --pal2019 --aa_pdb 1S40.pdb --Ka_nucl 80
~~~

[Example 12]

The capsid assembly preset supports 2-bead amino-acids and 3-bead nucleotides, adds sequence-based Debye-Hückel electrostatics and also adds equivalent contacts for protein-protein and protein-RNA/DNA interfaces. Using the capsid preset, example 9a can be reproduced using:

~~~
       python submit.py --capsid --aa_pdb AB.pdb CC.pdb gRNA.pdb
            --nmol 60 30 1 --cmap_i inter.contacts
~~~

[Example 13]

To change parameters, such as to set the monovalent ion concentration to 0.01 M for calculating Debye length, the additional flag --iconc with the argument 0.01 can be given:

~~~
       python submit.py --capsid --aa_pdb AB.pdb CC.pdb gRNA.pdb
            --nmol 60 30 1 --cmap_i inter.contacts --iconc 0.01
~~~

[Example 14]

For some models, the contact potential function can be modified. For example, instead of the default LJ-10-12 potential of the Cα-SBM^66^, a one minimum Gaussian pair potential^72^ can be used:

~~~
        python submit.py --calpha_sbm --aa_pdb 2CI2.pdb --contfunc 6
~~~

[Example 15]

Or with two minima Gaussian pair potential:

~~~
       python submit.py --calpha_sbm --aa_pdb 2CI2.pdb --contfunc 7
                        --cmap 2CI2_CA.contact
~~~

[Example 16]

As stated earlier, the two-minima Gaussian pair potential is only supported with a user defined contact map file (here 2CI2_CA.contact) with up to two native distance values.

For a given model, --cg_pdb can also be used instead of --aa_pdb, if the contact map is being calculated from the coarse-grained structure or is taken from the user-defined input (--cutofftype 2 or 0). Examples of preset models which take such contact maps are the SOP/SOP-SC models and their variants^44,67,69,88,89^. An example command for generating files for Protein L (PDB 1HZ6) with an SOP-SC model^44^ (flags: --maity2016 or --sopsc2016) is as follows:

~~~
           python submit.py --sopsc2016 --cg_pdb 1HZ6.CACB.pdb
~~~

[Example 17]

SuBMIT also supports the completely sequence-based potential, SOP-IDP^86,87^. This can be used without a native structure by using the --idp_seq flag and an input sequence. For example, to generate files for a 32-residue polyglutamine chain with SOP-IDP (--baidya2022 or -- sop_idp2022), the following command can be used:

~~~
            python submit.py --baidya2022 --idp_seq polyQ.fa
~~~

[Example 18]

An example of the input sequence file (here polyQ.fa) format is shown in Example 19b. While using SOP-IDP, the program generates a stretched chain coarse-grained structure file (.gro). If an input sequence flag (--idp_seq) is used along with an input structure and the SOP-SC (--sopsc2016) flag, then SuBMIT generates files for a hybrid SOP-SC-IDP (SOP-SC-IDR hybrid) model, where the regions indicated in the sequence file are taken to be intrinsically disordered regions (IDR) and are modelled using the SOP-IDP model^87^. The remaining regions are modelled using the structure-based 2016 SOP-SC model^44^. The SOP-IDP potential is also used to define interactions between the IDRs and the structured regions. This implementation is similar to a recently developed SOP-MULTI^88^ model. For both SOP-SC hybrid (--sopsc_idr or --sopsc2016 with --idr_seq) and SOP-MULTI (--baratam2024 or --sop_multi), an all-atom or coarse-grained input structure is necessary, along with the sequence file. The SuBMIT command is as follows:

~~~
python submit.py --sopsc_idr --aa_pdb 3HHR_B.pdb --idp_seq missing.fa
~~~

[Example 19a]

The file “missing.fa” (a similar format should be used for polyQ.fa) includes sequence, chain-id and residue number range of missing or disordered regions in the following format.

~~~
       >hGHR-ECD_Nterm:B:1:31
       FSGSEATAAILSRAPWSLQSVNPGLKTNSSK
       >hGHR-ECD_missing1:B:56:61
       HGTKNL
       >hGHR-ECD_missing1:B:73:74
       TQ
       >hGHR-ECD_Cterm:B:235:244
       QMSQFTCEED
~~~

[Example 19b]

SuBMIT also enables the use of compatible preset models to construct multimeric systems with symmetrized or equivalent contacts. For example, the SOP-SC and SOP-SC hybrid models can be implemented for a P53-tetramer with the N-terminal tail marked as a disordered region.

~~~
        python submit.py --aa_pdb 3TS8_A.pdb --sopsc_idr --nmol 4
              --idp_seq Nterminal.fa --cmap_i inter.contacts
~~~

[Example 20]

This command generates a 2-bead protein structure and topology file for a single P53 molecule, along with a combined topology file for four molecules. For both SOP-SC-IDR and SOP-MULTI, SuBMIT generates a fully extended coarse-grained structure file. Alternatively, the user can provide the full P53 monomer structure (after modeling missing regions using tools such as MODELLER^55^, AlphaFold^90^, ESMFold^91^, etc.) to the program along with the model preset and -- gen_cg flag, for generating a starting coarse-grained structure. A short MD run with the coarse-grained potential is recommended for getting an energy-minimized monomer structure, followed by the use of the GROMACS built-in genbox command to generate a structure with four P53 molecules.

## COMPARISON WITH PREVIOUS SIMULATIONS

We performed test simulations with most preset models that are present in the SuBMIT toolkit and the results were compared with previously published data. Here, we describe three such test cases. The first one describes folding of a single chain globular protein using the SOP-SC model^44^. The other two cases use a common Cα-SBM^66^: The second test case is simulations of the assembly of two copies of the same protein with symmetrized contacts (3D-domain-swapping)^45^. The third test case is the simulation of multiple copies of several protein fragments and how they interact with the full-length protein during folding. This involves equivalent interface contacts. All input files for the three test cases, including GROMACS parameter (.mdp) and OpenSMOG simulation input (.py) files are present in the example directory of the SuBMIT repository. Simulations were performed using OpenSMOG v1.1.1^43^ at the previously determined respective folding temperatures.

### Folding of Protein L using the SOP-SC model

The SOP-SC potential energy includes multiple functions that are not part of common MD packages. The original simulations of this system (Protein L)^44^ were implemented with “in-house” molecular dynamics code. To reproduce the data, we used SuBMIT to generate Protein L SOP-SC input files (with identical input parameters) for both GROMACS v4.5.4^40^ and OpenSMOG v1.1.1^43^ (with OpenMM 8.1.1^42^ API). We first preprocessed the Protein L all-atom structure (PDB: 1HZ6) by (i) removing chain B and C from the asymmetric unit, (ii) removing three N-terminal residues (HHA) and then (iii) adding hydrogen atoms to the structure using the UCSF-Chimera package^56^. The third step is needed primarily because the specific SOP-SC model^44^ assigns a sidechain (Cβ) bead for the Glycine residues at the position of the Cα H-atom (corresponding to a hypothetical L-Gly). It is important to note that SOP-SC models use kcal mol^-1^ as the default energy unit. For simulations in GROMACS or OpenSMOG, SuBMIT uses bonded and nonbonded interaction strengths converted into kJ mol^-1^ (values multiplied by 4.184 J cal^-1^). Since the Debye-length (Debye-Hückel electrostatics) depends on the (implicit) solution temperature, files were generated separately for each temperature run with the --debye_temp flag. For example,

~~~
python submit.py --sopsc2016 --aa_pdb 1HZ6_A_H.pdb --debye_temp 374.5
~~~

[Example 21]

Langevin dynamics (stochastic dynamics in GROMACS) simulations were performed at temperatures close to the published melting/folding temperature^44^, with replicates starting from both initial folded and unfolded states (each replicate run for 1E+9 steps). Trajectories were analyzed by calculating the radius of gyration (*R*_*g*_) of the chain and by using the number of formed native contacts to calculate the structure overlap function^44,92^, *χ*, where 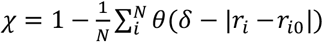. Here, N is the number of native contacts in the input structure and θ(x) is a step function with θ(x) = 0 if x < 0 and θ(x) = 1 otherwise. A native contact is formed if two atoms are part of the (native) contact list (*i*^*th*^contact pair of total N contacts) and the distance between them (*r*_*i*_) is within a cutoff (*δ* = 2Å) of the native distance *r*_*i*0_. Using this, structures in the folded ensemble have *χ* close to 0, while unfolded structures have *χ* close to 1. Figure 3A shows a 2-dimensional free energy profile (2D-FEP) which is a plot of the negative logarithm of the population of structures binned along *χ* (Y-axis) and *R*_*g*_ (X-axis), i.e., of *F* = −ln (*P*(*R*_*g*_, *χ*)). The calculated free energy profile (Figure 3A) and the folding pathways (average contact maps along *χ*, not shown) are in accordance with each other and with previous simulations^44^.

**Figure 3.**
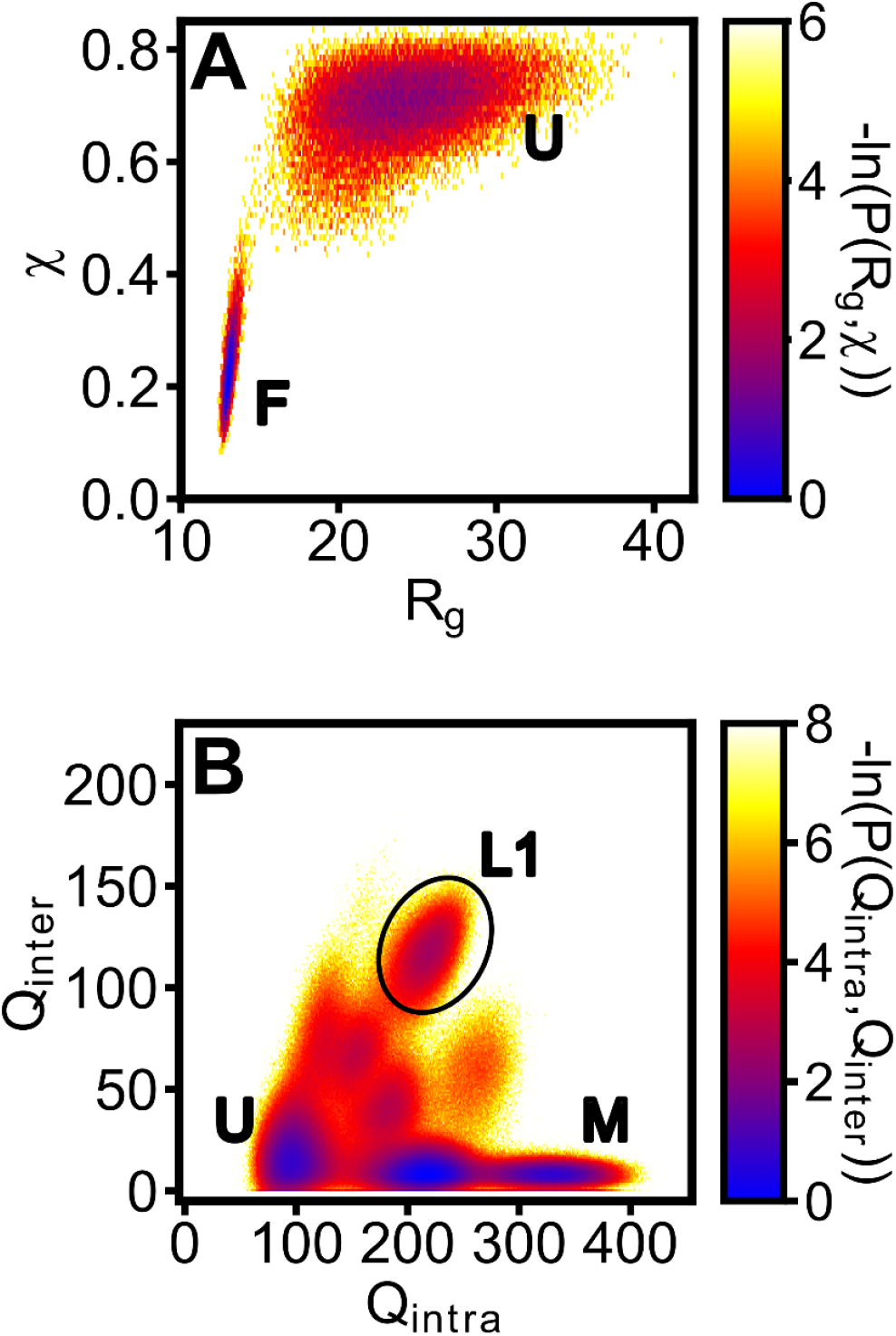
(A) 2-dimensional free energy profile (2D-FEP) for Protein L folding simulations using the SOP-SC model: Negative log of population of structures (free energies) with a given *χ* (structure-overlap function) and *R*_*g*_ (radius of gyration). The color bar is shown on the right. The unfolded (U) and folded (F) ensembles are marked. Results to be compared with Figure S2C from an earlier publication^44^. (B) 2D-FEP for SARS M^pro^ CTD domain swapping simulations using a common Cα-SBM. Negative log of population of structures with a given number of formed native interchain (*Q*_*inter*_) and intrachain (*Q*_*intra*_) contacts. The color bar is shown on the right. The unfolded (U), two-monomers folded (M) and the loop1 domain swapped (L1) states are marked. Results to be compared with Figure 2C of a previous publication^45^. Free energy baselines were adjusted to set the minimum value of both plots to 0.

### 3D-domain-swapping of the C-terminal domain of the SARS-CoV main protease (SARS M^pro^ CTD)

These simulations were originally performed^45^ using a common Cα-SBM^66^ which can be accessed in SuBMIT with the --calpha_sbm flag. For the comparison simulations, the structure of SARS M^pro^ CTD (residues 187-306, PDB: 2H2Z) was used to calculate a weighted contact map (--W_cont flag) and symmetrized using the --dswap flag as follows:

~~~
        python submit.py --calpha_sbm --aa_pdb 2H2Z_CTD.pdb --W_cont --dswap
~~~

[Example 22]

Two protein monomers were simulated, with a harmonic potential between their centroids preventing them from drifting apart. This option is supported in GROMACS (using “pull-code” options in the .mdp file). This option is not directly supported in OpenSMOG (v1.1.1) but can be implemented by using the OpenMM CustomCentroidBondForce() function in the OpenSMOG simulation input python script. Alternatively, it can also be implemented using the OpenSMOGmod plugin (https://github.com/sglabncbs/OpenSMOGmod) which supports adding energy functions between the center of mass of two (or more) chains in the OpenSMOG XML (.xml) file. Identical parameters were used with GROMACS and OpenSMOG, with multiple Langevin dynamics simulations performed at the folding temperature (determined in the previous study^45^) starting from both folded and unfolded initial structures (each for 2E+9 steps). The number of formed native interchain (*Q*_*inter*_) and intrachain (*Q*_*intra*_) contacts were calculated for each snapshot in the trajectory to identify the “folded-ness” of the protein monomers or dimer. A native contact is formed in a snapshot if the two atoms are part of the native contact list and are within 1.2 times the corresponding native distance in the monomer structure. Figure 3B shows a 2D-free energy profile, a plot of the negative logarithm of the population of structures binned along *Q*_*inter*_(Y-axis) and *Q*_*intra*_(X-axis), i.e. of, *F* = −ln (*P*(*Q*_*intra*_, *Q*_*inter*_). The 2D-FEP (Figure 3B) and the folding pathways (average contact maps, not shown) are in agreement with each other and with previous simulations^45^.

### Folding of SARS-CoV-2 M^pro^ C-terminal domain (SARS2 M^pro^ CTD) in the presence of self-peptides

In previous work^46^, the folding of SARS2 M^pro^ CTD was simulated in the presence of single self-peptides (a self-peptide is a segment of SARS2 M^pro^ CTD, termed the parent protein; six different self-peptides were simulated separately). A harmonic constraint between the self-peptide and the parent protein kept the two from drifting apart and was also used to mimic the effect of peptide concentration. In SuBMIT test simulations, the protocol was modified to include a periodic box with a single copy of SARS2 M^pro^ CTD along with 200 copies of each of the six self-peptides (a total of 200 × 6 = 1200 peptides; see Table 3 and Video 1). As done in the previous study^46^, the strength of the intra-chain self-peptide (intra-peptide) contacts was increased by five to mimic a pre-folded (or “stapled”) peptide. No additional harmonic constraints were used between the peptides and the SARS2 M^pro^ CTD. The input files were generated as follows:

**Table 3.**
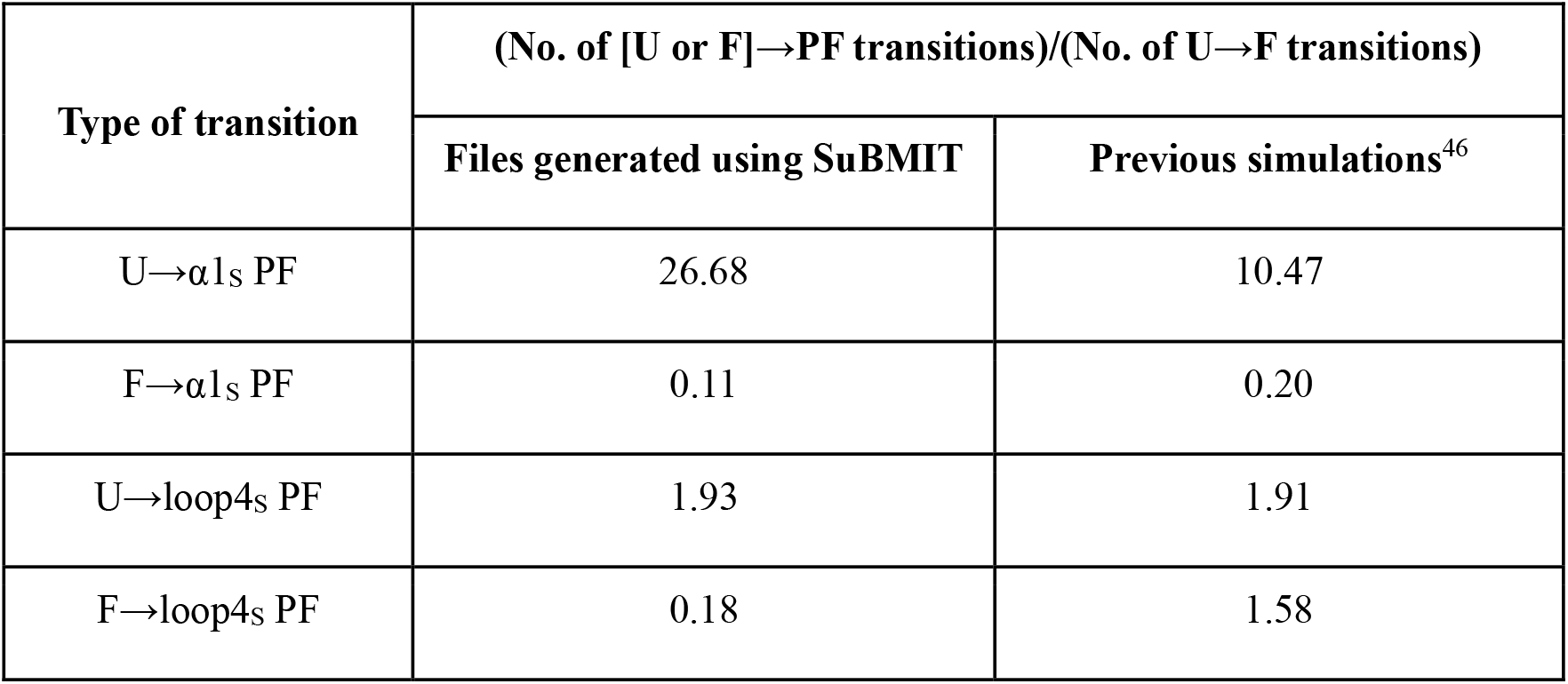
Results of SARS2 M^pro^ CTD folding simulations in the presence of its self-peptides. Previous simulations^46^ (right column) showed that the PF ensemble (ensemble in which the self-peptide replaces the native segment from the folded protein) is significantly populated only with the α1_S_ and the loop4_S_ self-peptides.

~~~
          python submit.py --aa_pdb 6Y2E_CTD.pdb prot1.pdb prot2.pdb
          prot3.pdb prot4.pdb prot5.pdb prot6.pdb --cmap 6Y2E_CTD.cont
          stap_prot1.cont stap_prot2.cont stap_prot3.cont stap_prot4.cont
          stap_prot5.cont stap_prot6.cont --cmap_i inter.cont --nmol 1 200
          200 200 200 200 200 --calpha_sbm --opensmog
~~~

[Example 23]

Here, 6Y2E_CTD.pdb (PDB ID: 6Y2E, residues 187-306) and 6Y2E_CTD.cont are the PDB and the Cα contact list files for SARS2 M^pro^ CTD. The protX.pdb and stap_protX.cont (where X is 1, 2, 3, 4, 5 or 6) are the PDB and Cα contact files for the six self-peptides from the SARS2 M^pro^ CTD structure, namely, (1) α1_S_, (2) α2_S_, (3) α3_S_, (4) α4_S_, (5) α5_S_ and (6) loop4_S_ where the “s” stands for stabilized. The inter.cont files include all the interchain contacts whose format is described in the earlier “**Contact-maps:**” subsection. For the parent protein (SARS2 M^pro^ CTD; given as the first input), prot0 was used as the chain identifier with prot1 to prot6 used for the six peptides. Since multiple copies of peptides were added to the system, SuBMIT adds the interchain contacts as part of the nonbonded parameters (equivalent contacts). Two simulation replicates (each of 2E+9 steps simulated using Langevin dynamics) were performed using OpenSMOG v1.1.1 with a 500 x 500 x 500 Å^3^ periodic box. The simulation temperature (0.99 k_B_T; k_B_ = Boltzmann constant) and all the other parameters were identical to those used in the original study^46^.

The number of native intrachain protein (M^pro^ CTD) contacts (*Q*_*intra*_) formed and the number of native interchain protein-peptide contacts (*Q*_*inter*_) formed were calculated for each simulation snapshot. A native contact is formed if the two atoms are part of the input contact list and are within 1.2 times the corresponding native distance in the snapshot. The number of formed interchain protein-peptide contacts 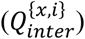 were determined separately for each of the 200 copies (indexed with *i*) of each of the six self-peptide types (*x* ∈ {1,2,3,4,5,6}). The *Q*_*intra*_ and the 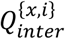 parameters were used to count the number of one-way transitions between the unfolded (U), folded (F) and the peptide-folded (PF) ensembles (a minimum where the self-peptide displaces its equivalent segment from the parent protein and they fold together into a single structure). For each peptide type (*x*), the 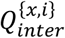 threshold for identifying the peptide-folded state, was the same as used in the previous study^46^. Despite using a system where each peptide can compete with all other peptides for binding to the folding protein, our simulations show, in agreement with previous results^46^, that only α1_S_ and loop4_S_ self-peptides lead to the formation of peptide-folded states. Moreover, the relative number of one-way transitions (see Table 3) for the two “inserting” peptide types, were comparable to the previously shown results involving a single peptide copy^46^. Interestingly, as compared to the previous study, the current approach shows a lower percentage of transitions from the folded to the peptide-folded (F→PF) states. In the original study, the higher number may be attributed to the presence of the harmonic constraint between the protein and the self-peptide, which can keep the self-peptide near the protein even when it is fully folded.

## COMPARISON WITH OTHER PACKAGES WHICH GENERATE SBM INPUT FILES

The overall goal and functionality of SuBMIT is similar to previously developed programs which generate SBM input files such as SMOG2^43,63^, Go-Kit^64^, eSBMtools^93,94^, SBMOpenMM^95^ and GoCα^25^. However, the primary purpose and scope of each of these packages is different. We compare and contrast SuBMIT with these programs.

SuBMIT generates input files for GROMACS and OpenMM/OpenSMOG and has been specifically developed for coarse-grained models, providing diverse preset protein and nucleic acid CG-SBMs. But it lacks support for all-atom models. Files for all-atom SBMs^60^ for both proteins and nucleic acids can be generated using the SMOG2^43,63^ package (and the SMOG-server webtool which was the earliest package developed). Packages such as eSBMtools^93^ and SBMOpenMM^95^, can also generate all-atom SBM files, although SBMOpenMM doesn’t include support for nucleic acid models.

Not many other protein CG-SBMs besides the Cα-SBM^66^ are included with the default installations of SMOG2, eSBMtools or SBMOpenMM. However, support for new CG-SBMs can be added to both SMOG2 and eSBMtools by creating model specific-files. Go-Kit^64^ offers support for both Cα-Cβ^96^ and Cα-SBMs. Both Go-Kit and eSBMtools also provide support for CG-SBMs with non-native sequence-based (hydrophobic or electrostatics) interactions. The GoCα web-tool^25^ is specifically designed for generating Cα SBM topology files with equivalent contacts, to study protein assembly and currently lacks support for all other (all-atom or CG) SBMs.

Unlike the other tools, SuBMIT provides support for some nucleic acid CG-SBMs. SuBMIT also provides input files for structure-independent protein-protein and protein-nucleotide interactions (electrostatics, hydrophobic and IDP potentials). These features together with the support for equivalent and symmetric contacts enables SuBMIT to generate input files for simulating the multimeric folding (such as 3D-domain swapping) and assembly (such as of protein-nucleic acid complexes) of varied biomolecular complexes using multiple variant CG-SBMs. Additionally, like SMOG2 and SBMOpenMM, SuBMIT can also encode single-minimum or dual-minima Gaussian functions (Figure 2). These are useful for encoding non-native attractive potentials or simulating conformational transitions with dual-minimum contacts.

In order to make SBMs more accessible, all the tools (with the exception of SBMOpenMM) generate input files for GROMACS, which is an open-source software with active development, and CPU parallelization support. However, custom bonded and non-bonded potential energy functions (like those used in many CG-SBMs) can only be implemented in GROMACS using a user-defined look-up table. GROMACS does not support parallelization on graphics processing units (GPU) while using look-up tables, limiting the calculation speed and resultant use of custom functions in larger biomolecular systems. For instance, the LJ-10-12 contact potential (Figure 2), which is the default in a common Cα-SBM^66^, can only be implemented using a user-defined look-up table in GROMACS. So, the GoCα web-tool (which is targeted towards studying protein assembly) replaces the LJ-10-12 with the LJ-6-12 (Figure 2) contact potential which is implemented in GROMACS and which can be used without look-up tables. OpenMM^41,42^ overcomes this limitation of GROMACS, by allowing direct implementation of, and GPU support for, custom/non-standard potential energy functions. This makes OpenMM more suitable for use with CG-SBMs. SuBMIT, SMOG2 and SBMOpenMM support generating files for OpenMM. In fact, SBMOpenMM is vertically integrated with OpenMM and doesn’t generate separate topology files. It uses the input protein structure to encode either all-atom or Cα-SBM potentials and directly executes the simulation using OpenMM libraries. This makes the process straightforward but restricts the user to pre-encoded models. Unlike SBMOpenMM, both SMOG2 and SuBMIT, first generate structure and SBM topology files, which can then be used with OpenSMOG, a tool specifically built for performing SBM simulations using the OpenMM API. This two-step process makes it easier for SuBMIT to support diverse CG-SBM flavors.

Overall SuBMIT is useful because it includes support for (a) diverse preset protein CG-SBMs, (b) CG-SBMs of nucleic acids, (c) custom energy functions and structure-independent interactions and (d) large multimeric systems. SuBMIT is modular in its implementation which allows incorporation of new model presets without altering the predefined classes and functions in the program. Thus, new models can be easily implemented and tested within SuBMIT, which if deposited to the SuBMIT repository, will make them accessible for use by all.

## CONCLUSIONS

There are currently several MD packages optimized for biomolecular simulations such as GROMACS and OpenMM. These have specific in-built potential energy forms which together are used to implement standard biomolecular forcefields such as AMBER and CHARMM. SuBMIT is a software package which uses these same potential energy forms to provide input files for performing MD simulations of coarse-grained structure-based models (CG-SBMs) of proteins, nucleic acids and a combination of the two. Additionally, support for OpenSMOG, an extension of OpenMM for SBMs, allows users to implement custom nonbonded and pair potential functions. SuBMIT is especially useful for generating simulation input files for large multimeric systems with multiple copies of different molecules. Being an open-source tool, the SuBMIT code can be downloaded and modified to add new models, which, if required, can be integrated into the main SuBMIT package. Overall, implementing CG-SBMs using SuBMIT allows the simulation of both pre-tested and new bespoke coarse-grained models on MD packages that are already optimized for parallelization on both CPU and GPU.

## Conflict of interest

There is no conflict of interest.

## Acknowledgements

We thank Dr. Hiranmay Maity. Dr. Lipika Baidya and Prof. Govardhan Reddy, SSCU, IISc, Bangalore, India for providing files for encoding and testing SOPSC and SOP-IDP models. We thank Dr. Sridhar Neelamraju and Dr. Sahil Lall for helpful discussions and Dr. Aravind Ravichandran, Antony John and Naren C for helping test SUBMIT. A part of the testing simulations was performed on PARAM Brahma HPC at IISER, Pune, India with access provided through the National Supercomputing Mission (NSM) grant MeitY/R\&D/HPC/2(1)/2014. This work was supported in part by the Department of Atomic Energy, Government of India through the Tata Institute of Fundamental Research Project Identification No. RTI 4006, the Simons Foundation (Grant No. 287975) and SERB Grant: CRG/2021/004754. DLP was supported by a SRF from Council of Scientific & Industrial Research (CSIR), India.

## Notes

https://github.com/sglabncbs/submit

### Competing Interest Statement

The authors have declared no competing interest.

## REFERENCES

(1) Pak, A. J.; Voth, G. A. Advances in Coarse-Grained Modeling of Macromolecular Complexes. Curr Opin Struct Biol 2018, 52, 119–126. 10.1016/j.sbi.2018.11.005.

(2) Singh, N.; Li, W. Recent Advances in Coarse-Grained Models for Biomolecules and Their Applications. Int J Mol Sci 2019, 20 (15), 3774. 10.3390/ijms20153774.

(3) Takada, S. Coarse-Grained Molecular Simulations of Large Biomolecules. Current Opinion in Structural Biology 2012, 22 (2), 130–137. 10.1016/j.sbi.2012.01.010.

(4) Takada, S.; Kanada, R.; Tan, C.; Terakawa, T.; Li, W.; Kenzaki, H. Modeling Structural Dynamics of Biomolecular Complexes by Coarse-Grained Molecular Simulations. Acc. Chem. Res. 2015, 48 (12), 3026–3035. 10.1021/acs.accounts.5b00338.

(5) Onuchic, J. N.; Luthey-Schulten, Z.; Wolynes, P. G. THEORY OF PROTEIN FOLDING: The Energy Landscape Perspective. Annual Review of Physical Chemistry 1997, 48 (1), 545–600. 10.1146/annurev.physchem.48.1.545.

(6) Onuchic, J. N.; Wolynes, P. G. Theory of Protein Folding. Current Opinion in Structural Biology 2004, 14 (1), 70–75. 10.1016/j.sbi.2004.01.009.

(7) Taketomi, H.; Ueda, Y.; Gō, N. Studies on Protein Folding, Unfolding and Fluctuations by Computer Simulation. International Journal of Peptide and Protein Research 1975, 7 (6), 445–459. 10.1111/j.1399-3011.1975.tb02465.x.

(8) Hills, R.; Brooks, C. Insights from Coarse-Grained Gō Models for Protein Folding and Dynamics. International Journal of Molecular Sciences 2009, 10 (3), 889–905. 10.3390/ijms10030889.

(9) Mascarenhas, N. M.; Gosavi, S. Understanding Protein Domain-Swapping Using Structure-Based Models of Protein Folding. Prog Biophys Mol Biol 2017, 128, 113–120. 10.1016/j.pbiomolbio.2016.09.013.

(10) Sinner, C.; Lutz, B.; John, S.; Reinartz, I.; Verma, A.; Schug, A. Simulating Biomolecular Folding and Function by Native-Structure-Based/Go-Type Models. Israel Journal of Chemistry 2014, 54 (8–9), 1165–1175. 10.1002/ijch.201400012.

(11) Takada, S. Gō Model Revisited. Biophysics and Physicobiology 2019, 16, 248–255. 10.2142/biophysico.16.0_248.

(12) Yadahalli, S.; Hemanth Giri Rao, V. V.; Gosavi, S. Modeling Non-Native Interactions in Designed Proteins. Israel Journal of Chemistry 2014. 10.1002/ijch.201400035.

(13) Thirumalai, D.; Hyeon, C. RNA and Protein Folding: Common Themes and Variations. Biochemistry 2005, 44 (13), 4957–4970. 10.1021/bi047314+.

(14) Hyeon, C.; Thirumalai, D. Mechanical Unfolding of RNA Hairpins. Proceedings of the National Academy of Sciences 2005, 102 (19), 6789–6794. 10.1073/pnas.0408314102.

(15) Jonikas, M. A.; Radmer, R. J.; Laederach, A.; Das, R.; Pearlman, S.; Herschlag, D.; Altman, R. B. Coarse-Grained Modeling of Large RNA Molecules with Knowledge-Based Potentials and Structural Filters. RNA 2009, 15 (2), 189–199. 10.1261/rna.1270809.

(16) Denesyuk, N. A.; Thirumalai, D. Coarse-Grained Model for Predicting RNA Folding Thermodynamics. Journal of Physical Chemistry B 2013. 10.1021/jp401087x.

(17) Poma, A. B.; Cieplak, M.; Theodorakis, P. E. Combining the MARTINI and Structure-Based Coarse-Grained Approaches for the Molecular Dynamics Studies of Conformational Transitions in Proteins. J. Chem. Theory Comput. 2017, 13 (3), 1366–1374. 10.1021/acs.jctc.6b00986.

(18) Mahmood, M. I.; Poma, A. B.; Okazaki, K. Optimizing Gō-MARTINI Coarse-Grained Model for F-BAR Protein on Lipid Membrane. Front. Mol. Biosci. 2021, 8. 10.3389/fmolb.2021.619381.

(19) Yang, S.; Song, C. Switching Gō-Martini for Investigating Protein Conformational Transitions and Associated Protein–Lipid Interactions. J. Chem. Theory Comput. 2024, 20 (6), 2618–2629. 10.1021/acs.jctc.3c01222.

(20) Giri Rao, V. V. H.; Desikan, R.; Ayappa, K. G.; Gosavi, S. Capturing the Membrane-Triggered Conformational Transition of an α-Helical Pore-Forming Toxin. J. Phys. Chem. B 2016, 120 (47), 12064–12078. 10.1021/acs.jpcb.6b09400.

(21) Shental-Bechor, D.; Smith, M. T. J.; MacKenzie, D.; Broom, A.; Marcovitz, A.; Ghashut, F.; Go, C.; Bralha, F.; Meiering, E. M.; Levy, Y. Nonnative Interactions Regulate Folding and Switching of Myristoylated Protein. Proceedings of the National Academy of Sciences 2012, 109 (44), 17839–17844. 10.1073/pnas.1201803109.

(22) Levy, Y.; Wolynes, P. G.; Onuchic, J. N. Protein Topology Determines Binding Mechanism. Proceedings of the National Academy of Sciences 2004, 101 (2), 511–516. 10.1073/pnas.2534828100.

(23) Levy, Y.; Onuchic, J. N. Mechanisms of Protein Assembly: Lessons from Minimalist Models. Acc. Chem. Res. 2006, 39 (2), 135–142. 10.1021/ar040204a.

(24) Levy, Y.; Papoian, G. A.; Onuchic, J. N.; Wolynes, P. G. Energy Landscape Analysis of Protein Dimers. Israel Journal of Chemistry 2004, 44 (1–3), 281–297. 10.1560/XGCB-WUHN-JRJC-BV0Y.

(25) Walter, L. J.; Quoika, P. K.; Zacharias, M. Structure-Based Protein Assembly Simulations Including Various Binding Sites and Conformations. J. Chem. Inf. Model. 2024, 64 (8), 3465–3476. 10.1021/acs.jcim.4c00212.

(26) Scalone, E.; Broggini, L.; Visentin, C.; Erba, D.; Bačić Toplek, F.; Peqini, K.; Pellegrino, S.; Ricagno, S.; Paissoni, C.; Camilloni, C. Multi-eGO: An in Silico Lens to Look into Protein Aggregation Kinetics at Atomic Resolution. Proceedings of the National Academy of Sciences 2022, 119 (26), e2203181119. 10.1073/pnas.2203181119.

(27) Bačić Toplek, F.; Scalone, E.; Stegani, B.; Paissoni, C.; Capelli, R.; Camilloni, C. Multi-eGO: Model Improvements toward the Study of Complex Self-Assembly Processes. J Chem Theory Comput 2024, 20 (1), 459–468. 10.1021/acs.jctc.3c01182.

(28) Givaty, O.; Levy, Y. Protein Sliding along DNA: Dynamics and Structural Characterization. Journal of Molecular Biology 2009, 385 (4), 1087–1097. 10.1016/j.jmb.2008.11.016.

(29) Mishra, G.; Levy, Y. Molecular Determinants of the Interactions between Proteins and ssDNA. Proceedings of the National Academy of Sciences of the United States of America 2015. 10.1073/pnas.1416355112.

(30) Pal, A.; Levy, Y. Structure, Stability and Specificity of the Binding of ssDNA and ssRNA with Proteins. PLoS Computational Biology 2019. 10.1371/journal.pcbi.1006768.

(31) Baweja, L.; Roche, J. Pushing the Limits of Structure-Based Models: Prediction of Nonglobular Protein Folding and Fibrils Formation with Go-Model Simulations. J Phys Chem B 2018, 122 (9), 2525–2535. 10.1021/acs.jpcb.7b12129.

(32) Yang, S.; Cho, S. S.; Levy, Y.; Cheung, M. S.; Levine, H.; Wolynes, P. G.; Onuchic, J. N. Domain Swapping Is a Consequence of Minimal Frustration. Proceedings of the National Academy of Sciences of the United States of America 2004, 101 (38), 13786–13791. 10.1073/pnas.0403724101.

(33) Yang, S.; Levine, H.; Onuchic, J. N. Protein Oligomerization Through Domain Swapping: Role of Inter-Molecular Interactions and Protein Concentration. Journal of Molecular Biology 2005, 352 (1), 202–211. 10.1016/j.jmb.2005.06.062.

(34) Ding, F.; Prutzman, K. C.; Campbell, S. L.; Dokholyan, N. V. Topological Determinants of Protein Domain Swapping. Structure 2006, 14 (1), 5–14. 10.1016/j.str.2005.09.008.

(35) Nelson, M. T.; Humphrey, W.; Gursoy, A.; Dalke, A.; Kalé, L. V.; Skeel, R. D.; Schulten, K. NAMD: A Parallel, Object-Oriented Molecular Dynamics Program. The International Journal of Supercomputer Applications and High Performance Computing 1996, 10 (4), 251–268. 10.1177/109434209601000401.

(36) Phillips, J. C.; Hardy, D. J.; Maia, J. D. C.; Stone, J. E.; Ribeiro, J. V.; Bernardi, R. C.; Buch, R.; Fiorin, G.; Hénin, J.; Jiang, W.; McGreevy, R.; Melo, M. C. R.; Radak, B. K.; Skeel, R. D.; Singharoy, A.; Wang, Y.; Roux, B.; Aksimentiev, A.; Luthey-Schulten, Z.; Kalé, L. V.; Schulten, K.; Chipot, C.; Tajkhorshid, E. Scalable Molecular Dynamics on CPU and GPU Architectures with NAMD. The Journal of Chemical Physics 2020, 153 (4), 044130. 10.1063/5.0014475.

(37) Thompson, A. P.; Aktulga, H. M.; Berger, R.; Bolintineanu, D. S.; Brown, W. M.; Crozier, P. S.; in ‘t Veld, P. J.; Kohlmeyer, A.; Moore, S. G.; Nguyen, T. D.; Shan, R.; Stevens, M. J.; Tranchida, J.; Trott, C.; Plimpton, S. J. LAMMPS - a Flexible Simulation Tool for Particle-Based Materials Modeling at the Atomic, Meso, and Continuum Scales. Computer Physics Communications 2022, 271, 108171. 10.1016/j.cpc.2021.108171.

(38) Berendsen, H. J. C.; van der Spoel, D.; van Drunen, R. GROMACS: A Message-Passing Parallel Molecular Dynamics Implementation. Computer Physics Communications 1995, 91 (1), 43–56. 10.1016/0010-4655(95)00042-E.

(39) Van Der Spoel, D.; Lindahl, E.; Hess, B.; Groenhof, G.; Mark, A. E.; Berendsen, H. J. C. GROMACS: Fast, Flexible, and Free. Journal of Computational Chemistry 2005, 26 (16), 1701–1718. 10.1002/jcc.20291.

(40) Pronk, S.; Páll, S.; Schulz, R.; Larsson, P.; Bjelkmar, P.; Apostolov, R.; Shirts, M. R.; Smith, J. C.; Kasson, P. M.; van der Spoel, D.; Hess, B.; Lindahl, E. GROMACS 4.5: A High-Throughput and Highly Parallel Open Source Molecular Simulation Toolkit. Bioinformatics 2013, 29 (7), 845–854. 10.1093/bioinformatics/btt055.

(41) Eastman, P.; Swails, J.; Chodera, J. D.; McGibbon, R. T.; Zhao, Y.; Beauchamp, K. A.; Wang, L. P.; Simmonett, A. C.; Harrigan, M. P.; Stern, C. D.; Wiewiora, R. P.; Brooks, B. R.; Pande, V. S. OpenMM 7: Rapid Development of High Performance Algorithms for Molecular Dynamics. PLOS Computational Biology 2017, 13 (7), e1005659. 10.1371/JOURNAL.PCBI.1005659.

(42) Eastman, P.; Galvelis, R.; Peláez, R. P.; Abreu, C. R. A.; Farr, S. E.; Gallicchio, E.; Gorenko, A.; Henry, M. M.; Hu, F.; Huang, J.; Krämer, A.; Michel, J.; Mitchell, J. A.; Pande, V. S.; Rodrigues, J. P.; Rodriguez-Guerra, J.; Simmonett, A. C.; Singh, S.; Swails, J.; Turner, P.; Wang, Y.; Zhang, I.; Chodera, J. D.; De Fabritiis, G.; Markland, T. E. OpenMM 8: Molecular Dynamics Simulation with Machine Learning Potentials. J. Phys. Chem. B 2024, 128 (1), 109–116. 10.1021/acs.jpcb.3c06662.

(43) de Oliveira, A. B.; Contessoto, V. G.; Hassan, A.; Byju, S.; Wang, A.; Wang, Y.; Dodero-Rojas, E.; Mohanty, U.; Noel, J. K.; Onuchic, J. N.; Whitford, P. C. SMOG 2 and OpenSMOG: Extending the Limits of Structure-Based Models. Protein science : a publication of the Protein Society 2022, 31 (1), 158–172. 10.1002/PRO.4209.

(44) Maity, H.; Reddy, G. Folding of Protein L with Implications for Collapse in the Denatured State Ensemble. J. Am. Chem. Soc. 2016, 138 (8), 2609–2616. 10.1021/jacs.5b11300.

(45) Terse, V. L.; Gosavi, S. The Molecular Mechanism of Domain Swapping of the C-Terminal Domain of the SARS-Coronavirus Main Protease. Biophysical Journal 2021, 120 (3), 504–516. 10.1016/j.bpj.2020.11.2277.

(46) Banerjee, A.; Gosavi, S. Potential Self-Peptide Inhibitors of the SARS-CoV-2 Main Protease. J. Phys. Chem. B 2023, 127 (4), 855–865. 10.1021/acs.jpcb.2c05917.

(47) Huang, J.; Rauscher, S.; Nawrocki, G.; Ran, T.; Feig, M.; de Groot, B. L.; Grubmüller, H.; MacKerell, A. D. CHARMM36m: An Improved Force Field for Folded and Intrinsically Disordered Proteins. Nat Methods 2017, 14 (1), 71–73. 10.1038/nmeth.4067.

(48) Denning, E. J.; Priyakumar, U. D.; Nilsson, L.; Mackerell Jr., A. D. Impact of 2′-Hydroxyl Sampling on the Conformational Properties of RNA: Update of the CHARMM All-Atom Additive Force Field for RNA. Journal of Computational Chemistry 2011, 32 (9), 1929–1943. 10.1002/jcc.21777.

(49) Hart, K.; Foloppe, N.; Baker, C. M.; Denning, E. J.; Nilsson, L.; MacKerell, A. D. Jr. Optimization of the CHARMM Additive Force Field for DNA: Improved Treatment of the BI/BII Conformational Equilibrium. J. Chem. Theory Comput. 2012, 8 (1), 348–362. 10.1021/ct200723y.

(50) Lindorff-Larsen, K.; Piana, S.; Palmo, K.; Maragakis, P.; Klepeis, J. L.; Dror, R. O.; Shaw, D. E. Improved Side-Chain Torsion Potentials for the Amber ff99SB Protein Force Field. Proteins 2010, 78 (8), 1950–1958. 10.1002/prot.22711.

(51) Maier, J. A.; Martinez, C.; Kasavajhala, K.; Wickstrom, L.; Hauser, K. E.; Simmerling, C. ff14SB: Improving the Accuracy of Protein Side Chain and Backbone Parameters from ff99SB. J Chem Theory Comput 2015, 11 (8), 3696–3713. 10.1021/acs.jctc.5b00255.

(52) Tian, C.; Kasavajhala, K.; Belfon, K. A. A.; Raguette, L.; Huang, H.; Migues, A. N.; Bickel, J.; Wang, Y.; Pincay, J.; Wu, Q.; Simmerling, C. ff19SB: Amino-Acid-Specific Protein Backbone Parameters Trained against Quantum Mechanics Energy Surfaces in Solution. J. Chem. Theory Comput. 2020, 16 (1), 528–552. 10.1021/acs.jctc.9b00591.

(53) Grosse-Kunstleve, R. W. hybrid-36: PDB serial and sequence numbers. https://cci.lbl.gov/hybrid_36/.

(54) Grosse-Kunstleve, R. W.; Sauter, N. K.; Moriarty, N. W.; Adams, P. D. The Computational Crystallography Toolbox: Crystallographic Algorithms in a Reusable Software Framework. J Appl Cryst 2002, 35 (1), 126–136. 10.1107/S0021889801017824.

(55) Webb, B.; Sali, A. Comparative Protein Structure Modeling Using MODELLER. Current Protocols in Bioinformatics 2014, 47 (1), 5.6.1-5.6.32. 10.1002/0471250953.bi0506s47.

(56) Pettersen, E. F.; Goddard, T. D.; Huang, C. C.; Couch, G. S.; Greenblatt, D. M.; Meng, E. C.; Ferrin, T. E. UCSF Chimera - A Visualization System for Exploratory Research and Analysis. Journal of Computational Chemistry 2004, 25 (13), 1605–1612. 10.1002/jcc.20084.

(57) Noel, J. K.; Whitford, P. C.; Onuchic, J. N. The Shadow Map: A General Contact Definition for Capturing the Dynamics of Biomolecular Folding and Function. Journal of Physical Chemistry B 2012, 116 (29), 8692–8702. 10.1021/JP300852D/ASSET/IMAGES/MEDIUM/JP-2012-00852D_0011.GIF.

(58) Sobolev, V.; Sorokine, A.; Prilusky, J.; Abola, E. E.; Edelman, M. Automated Analysis of Interatomic Contacts in Proteins. Bioinformatics 1999, 15 (4), 327–332. 10.1093/bioinformatics/15.4.327.

(59) da Silveira, C. H.; Pires, D. E. V.; Minardi, R. C.; Ribeiro, C.; Veloso, C. J. M.; Lopes, J. C. D.; Meira, W.; Neshich, G.; Ramos, C. H. I.; Habesch, R.; Santoro, M. M. Protein Cutoff Scanning: A Comparative Analysis of Cutoff Dependent and Cutoff Free Methods for Prospecting Contacts in Proteins. Proteins 2009, 74 (3), 727–743. 10.1002/prot.22187.

(60) Whitford, P. C.; Noel, J. K.; Gosavi, S.; Schug, A.; Sanbonmatsu, K. Y.; Onuchic, J. N. An All-Atom Structure-Based Potential for Proteins: Bridging Minimal Models with All-Atom Empirical Forcefields. Proteins: Structure, Function, and Bioinformatics 2009, 75 (2), 430–441. 10.1002/prot.22253.

(61) Sugita, M.; Kikuchi, T. Incorporating into a Cα Go Model the Effects of Geometrical Restriction on Cα Atoms Caused by Side Chain Orientations. Proteins: Structure, Function, and Bioinformatics 2013, 81 (8), 1434–1445. 10.1002/prot.24294.

(62) Yadahalli, S.; Gosavi, S. Functionally Relevant Specific Packing Can Determine Protein Folding Routes. J Mol Biol 2016, 428 (2 Pt B), 509–521. 10.1016/j.jmb.2015.12.014.

(63) Noel, J. K.; Levi, M.; Raghunathan, M.; Lammert, H.; Hayes, R. L.; Onuchic, J. N.; Whitford, P. C. SMOG 2: A Versatile Software Package for Generating Structure-Based Models. PLOS Computational Biology 2016, 12 (3), e1004794. 10.1371/journal.pcbi.1004794.

(64) Neelamraju, S.; Wales, D. J.; Gosavi, S. Go-Kit: A Tool to Enable Energy Landscape Exploration of Proteins. Journal of Chemical Information and Modeling 2019, 59 (5), 1703–1708. 10.1021/acs.jcim.9b00007.

(65) Azia, A.; Levy, Y. Nonnative Electrostatic Interactions Can Modulate Protein Folding: Molecular Dynamics with a Grain of Salt. Journal of Molecular Biology 2009, 393 (2), 527–542. 10.1016/j.jmb.2009.08.010.

(66) Clementi, C.; Nymeyer, H.; Onuchic, J. N. Topological and Energetic Factors: What Determines the Structural Details of the Transition State Ensemble and “En-Route” Intermediates for Protein Folding? An Investigation for Small Globular Proteins. Journal of Molecular Biology 2000, 298 (5), 937–953. 10.1006/jmbi.2000.3693.

(67) Reddy, G.; Liu, Z.; Thirumalai, D. Denaturant-Dependent Folding of GFP. Proceedings of the National Academy of Sciences 2012, 109 (44), 17832–17838. 10.1073/pnas.1201808109.

(68) Wallin, S.; Chan, H. S. Conformational Entropic Barriers in Topology-Dependent Protein Folding: Perspectives from a Simple Native-Centric Polymer Model. J. Phys.: Condens. Matter 2006, 18 (14), S307. 10.1088/0953-8984/18/14/S14.

(69) Hyeon, C.; Dima, R. I.; Thirumalai, D. Pathways and Kinetic Barriers in Mechanical Unfolding and Refolding of RNA and Proteins. Structure 2006, 14 (11), 1633–1645. 10.1016/j.str.2006.09.002.

(70) Zarrine-Afsar, A.; Wallin, S.; Neculai, A. M.; Neudecker, P.; Howell, P. L.; Davidson, A. R.; Hue, S. C. Theoretical and Experimental Demonstration of the Importance of Specific Nonnative Interactions in Protein Folding. Proceedings of the National Academy of Sciences of the United States of America 2008. 10.1073/pnas.0801874105.

(71) Jayanthi, L. P.; Mascarenhas, N. M.; Gosavi, S. Structure Dictates the Mechanism of Ligand Recognition in the Histidine and Maltose Binding Proteins. Current Research in Structural Biology 2020, 2, 180–190. 10.1016/j.crstbi.2020.08.001.

(72) Lammert, H.; Schug, A.; Onuchic, J. N. Robustness and Generalization of Structure-Based Models for Protein Folding and Function. Proteins: Structure, Function, and Bioinformatics 2009, 77 (4), 881–891. 10.1002/prot.22511.

(73) Lu, Q.; Wang, J. Single Molecule Conformational Dynamics of Adenylate Kinase: Energy Landscape, Structural Correlations, and Transition State Ensembles. J Am Chem Soc 2008, 130 (14), 4772–4783. 10.1021/ja0780481.

(74) A, S.; Pc, W.; Y, L.; Jn, O. Mutations as Trapdoors to Two Competing Native Conformations of the Rop-Dimer. Proceedings of the National Academy of Sciences of the United States of America 2007, 104 (45). 10.1073/pnas.0706077104.

(75) Whitford, P. C.; Miyashita, O.; Levy, Y.; Onuchic, J. N. Conformational Transitions of Adenylate Kinase: Switching by Cracking. J Mol Biol 2007, 366 (5), 1661–1671. 10.1016/j.jmb.2006.11.085.

(76) Cheung, M. S.; García, A. E.; Onuchic, J. N. Protein Folding Mediated by Solvation: Water Expulsion and Formation of the Hydrophobic Core Occur after the Structural Collapse. Proceedings of the National Academy of Sciences of the United States of America 2002. 10.1073/pnas.022387699.

(77) Kaya, H.; Chan, H. S. Solvation Effects and Driving Forces for Protein Thermodynamic and Kinetic Cooperativity: How Adequate Is Native-Centric Topological Modeling? J Mol Biol 2003, 326 (3), 911–931. 10.1016/s0022-2836(02)01434-1.

(78) Liu, Z.; Chan, H. S. Solvation and Desolvation Effects in Protein Folding: Native Flexibility, Kinetic Cooperativity and Enthalpic Barriers under Isostability Conditions. Phys Biol 2005, 2 (4), S75–85. 10.1088/1478-3975/2/4/S01.

(79) Cho, S. S.; Levy, Y.; Onuchic, J. N.; Wolynes, P. G. Overcoming Residual Frustration in Domain-Swapping: The Roles of Disulfide Bonds in Dimerization and Aggregation. Physical biology 2005, 2 (2), S44–S55. 10.1088/1478-3975/2/2/S05.

(80) Mascarenhas, N. M.; Gosavi, S. Protein Domain-Swapping Can Be a Consequence of Functional Residues. Journal of Physical Chemistry B 2016, 120 (28), 6929–6938. 10.1021/acs.jpcb.6b03968.

(81) Das, A.; Yadav, A.; Gupta, M.; R, P.; Terse, V. L.; Vishvakarma, V.; Singh, S.; Nandi, T.; Banerjee, A.; Mandal, K.; Gosavi, S.; Das, R.; Ainavarapu, S. R. K.; Maiti, S. Rational Design of Protein-Specific Folding Modifiers. J. Am. Chem. Soc. 2021, 143 (44), 18766–18776. 10.1021/jacs.1c09611.

(82) Verlet, L. Computer “Experiments” on Classical Fluids. I. Thermodynamical Properties of Lennard-Jones Molecules. Phys. Rev. 1967, 159 (1), 98–103. 10.1103/PhysRev.159.98.

(83) Miyazawa, S.; Jernigan, R. L. Estimation of Effective Interresidue Contact Energies from Protein Crystal Structures: Quasi-Chemical Approximation. Macromolecules 1985, 18 (3), 534–552. 10.1021/ma00145a039.

(84) Miyazawa, S.; Jernigan, R. L. An Empirical Energy Potential with a Reference State for Protein Fold and Sequence Recognition. Proteins: Structure, Function and Genetics 1999. 10.1002/(SICI)1097-0134(19990815)36:3%253C357::AID-PROT10%253E3.0.CO;2-U.

(85) Betancourt, M. R.; Thirumalai, D. Pair Potentials for Protein Folding: Choice of Reference States and Sensitivity of Predicted Native States to Variations in the Interaction Schemes. Protein Sci 1999, 8 (2), 361–369. 10.1110/ps.8.2.361.

(86) Baul, U.; Chakraborty, D.; Mugnai, M. L.; Straub, J. E.; Thirumalai, D. Sequence Effects on Size, Shape, and Structural Heterogeneity in Intrinsically Disordered Proteins. J. Phys. Chem. B 2019, 123 (16), 3462–3474. 10.1021/acs.jpcb.9b02575.

(87) Baidya, L.; Reddy, G. pH Induced Switch in the Conformational Ensemble of Intrinsically Disordered Protein Prothymosin-α and Its Implications for Amyloid Fibril Formation. J. Phys. Chem. Lett. 2022, 13 (41), 9589–9598. 10.1021/acs.jpclett.2c01972.

(88) Baratam, K.; Srivastava, A. SOP-MULTI: A Self-Organized Polymer-Based Coarse-Grained Model for Multidomain and Intrinsically Disordered Proteins with Conformation Ensemble Consistent with Experimental Scattering Data. J. Chem. Theory Comput. 2024, 20 (22), 10179–10198. 10.1021/acs.jctc.4c00579.

(89) Reddy, G.; Thirumalai, D. Collapse Precedes Folding in Denaturant-Dependent Assembly of Ubiquitin. J Phys Chem B 2017, 121 (5), 995–1009. 10.1021/acs.jpcb.6b13100.

(90) Jumper, J.; Evans, R.; Pritzel, A.; Green, T.; Figurnov, M.; Ronneberger, O.; Tunyasuvunakool, K.; Bates, R.; Žídek, A.; Potapenko, A.; Bridgland, A.; Meyer, C.; Kohl, S. A. A.; Ballard, A. J.; Cowie, A.; Romera-Paredes, B.; Nikolov, S.; Jain, R.; Adler, J.; Back, T.; Petersen, S.; Reiman, D.; Clancy, E.; Zielinski, M.; Steinegger, M.; Pacholska, M.; Berghammer, T.; Bodenstein, S.; Silver, D.; Vinyals, O.; Senior, A. W.; Kavukcuoglu, K.; Kohli, P.; Hassabis, D. Highly Accurate Protein Structure Prediction with AlphaFold. Nature 2021, 596 (7873), 583–589. 10.1038/s41586-021-03819-2.

(91) Lin, Z.; Akin, H.; Rao, R.; Hie, B.; Zhu, Z.; Lu, W.; Smetanin, N.; Verkuil, R.; Kabeli, O.; Shmueli, Y.; Dos Santos Costa, A.; Fazel-Zarandi, M.; Sercu, T.; Candido, S.; Rives, A. Evolutionary-Scale Prediction of Atomic-Level Protein Structure with a Language Model. Science 2023, 379 (6637), 1123–1130. 10.1126/science.ade2574.

(92) Guo, Z.; Thirumalai, D. Kinetics and Thermodynamics of Folding of Ade NovoDesigned Four-Helix Bundle Protein. Journal of Molecular Biology 1996, 263 (2), 323–343. 10.1006/jmbi.1996.0578.

(93) Lutz, B.; Sinner, C.; Heuermann, G.; Verma, A.; Schug, A. eSBMTools 1.0: Enhanced Native Structure-Based Modeling Tools. Bioinformatics 2013, 29 (21), 2795–2796. 10.1093/bioinformatics/btt478.

(94) Lutz, B.; Sinner, C.; Bozic, S.; Kondov, I.; Schug, A. Native Structure-Based Modeling and Simulation of Biomolecular Systems per Mouse Click. BMC Bioinformatics 2014, 15 (1), 292. 10.1186/1471-2105-15-292.

(95) Floor, M.; Li, K.; Estévez-Gay, M.; Agulló, L.; Muñoz-Torres, P. M.; Hwang, J. K.; Osuna, S.; Villà-Freixa, J. SBMOpenMM: A Builder of Structure-Based Models for OpenMM. J. Chem. Inf. Model. 2021, 61 (7), 3166–3171. 10.1021/acs.jcim.1c00122.

(96) Cheung, M. S.; Finke, J. M.; Callahan, B.; Onuchic, J. N. Exploring the Interplay between Topology and Secondary Structural Formation in the Protein Folding Problem. J. Phys. Chem. B 2003, 107 (40), 11193–11200. 10.1021/jp034441r.

